# Genetic-Dependent Brain Signatures of Resilience: Interactions among Childhood Abuse, Genetic Risks and Brain Function

**DOI:** 10.1101/2024.09.16.612982

**Authors:** Han Lu, Edmund T. Rolls, Hanjia Liu, Dan J. Stein, Barbara J. Sahakian, Rebecca Elliott, Tianye Jia, Chao Xie, Shitong Xiang, Nan Wang, Tobias Banaschewski, Arun L.W. Bokde, Sylvane Desrivières, Herta Flor, Antoine Grigis, Hugh Garavan, Andreas Heinz, Rüdiger Brühl, Jean-Luc Martinot, Marie-Laure Paillère Martinot, Eric Artiges, Frauke Nees, Dimitri Papadopoulos Orfanos, Herve Lemaitre, Luise Poustka, Sarah Hohmann, Nathalie Holz, Juliane H. Fröhner, Michael N. Smolka, Nilakshi Vaidya, Henrik Walter, Robert Whelan, Gunter Schumann, Jianfeng Feng, Qiang Luo, the IMAGEN Consortium

## Abstract

Resilience to emotional disorders is critical for adolescent mental health, especially following childhood abuse. Yet, brain signatures of resilience remain undetermined due to the differential susceptibility of the brain’s emotion processing system to environmental stresses. Analyzing brain’s responses to angry faces in a longitudinally large-scale adolescent cohort (IMAGEN), we identified two functional networks related to the orbitofrontal and occipital regions as candidate brain signatures of resilience. In girls, but not boys, higher activation in the orbitofrontal-related network was associated with fewer emotional symptoms following childhood abuse, but only when the polygenic burden for depression was high. This finding defined a genetic-dependent brain (GDB) signature of resilience. Notably, this GDB signature predicted subsequent emotional disorders in late adolescence, extending into early adulthood and generalizable to another independent prospective cohort (ABCD). Our findings underscore the genetic modulation of resilience-brain connections, laying the foundation for enhancing adolescent mental health through resilience promotion.

## Introduction

Resilience, which is crucial for mental health, refers to the capacity for positive adaptation in coping with stress^1^. Childhood abuse (e.g., emotional abuse, physical abuse and sexual abuse), affecting over a billion people globally^2^, heightens the risk of emotional disorders such as depression and anxiety^3^. These disorders have been linked to dysfunctions in the brain’s emotion processing system (e.g., brain regions activated during emotion perception and emotion regulation)^4^, which is influenced by genetics during adolescent brain development^5,6^. Advanced knowledge of the genetic influences on resilience-brain associations can enhance prediction of emotional disorders following childhood abuse and aid in accurately identifying vulnerable individuals to facilitate early intervention.

In population-based neuroimaging studies, instead of categorizing resilient individuals from vulnerable ones, a neuroimaging marker (*i*.*e*., a brain signature) of resilience is often detected by an association where a higher level of this marker is associated with fewer emotional symptoms following childhood abuse^1,7^. This is highly relevant since there is an extensive literature showing that more emotional symptoms during childhood and adolescence are associated with higher risks (odds ratio = 1.85) of developing major depressive disorders during adulthood^8^. Previous studies of the brain’s signatures of resilience often focused on the fronto-limbic regions (e.g., the orbitofrontal cortex (OFC), medial prefrontal cortex, anterior cingulate cortex, amygdala, etc.)^9^. However, current findings in the literature are far from conclusive. For example, both hyper-^10^ and hypo-^11^ responses of the amygdala to negative emotional stimuli have been associated with fewer emotional symptoms following childhood abuse. Another example is that stronger spontaneous OFC activation has been associated with higher resilience as measured by the Connor-Davidson resilience scale in boys, but lower in girls ^12^. One source of these inconsistencies is that resilience can be built from optimized functions of various brain regions in different individuals as long as these optimizations can enhance the brain’s capability of emotion processing^13^. Therefore, instead of individual brain regions, the brain’s signatures of resilience might be better identified by the brain’s functional networks for emotion processing.

As hypothesized by the differential susceptibility theory ^14^, another source of these inconsistencies arises from the complex three-way interactions among childhood abuse, the brain’s emotion processing system, and genetic risk for depression. In the literature, various genetic variations in depression-related genes, such as 5HTTLPR^15^ and FKBP5^16^, interact with childhood maltreatment and alter the functional connectivity of the amygdala within the brain’s emotion circuit. A high polygenic risk score for major depressive disorder (PRS_MDD_) has been reported to interact with childhood trauma, increasing the susceptibility to developing more emotional symptoms^17^. Therefore, it is possible to detect a genetic-dependent brain signature of resilience (GBDSR) by a three-way interaction, where PRS_MDD_ modulates the association between a higher level of this brain signature and fewer emotional symptoms following childhood abuse. However, previously the understanding of the three-way interaction was limited, mainly due to the lack of neuroimaging data with a sufficiently large sample size activating the brain’s emotion processing system. The IMAGEN study, a large-scale neuroimaging cohort ^18^, used the emotional face task in a functional magnetic resonance imaging experiment to probe the brain’s emotion processing system^19^.

To address the above problems, we aim to answer the following four main questions regarding the brain signatures of resilience to developing more emotional symptoms following childhood abuse in the context of genetic predispositions for depression (Figure 1). (1) Can we isolate distinct functional networks in the brain’s emotion processing system as candidate signatures for resilience? (2) Can we identify the GDBSR by detecting significant three-way interactions among these functional networks, childhood abuse and PRS_MDD_ in relation to emotional symptoms? (3) To be clinically relevant, can these identified GDBSR predict subsequent emotional disorders following childhood abuse? (4) Are these predictions generalizable to other developmental stages and independent datasets?

**Figure 1.**
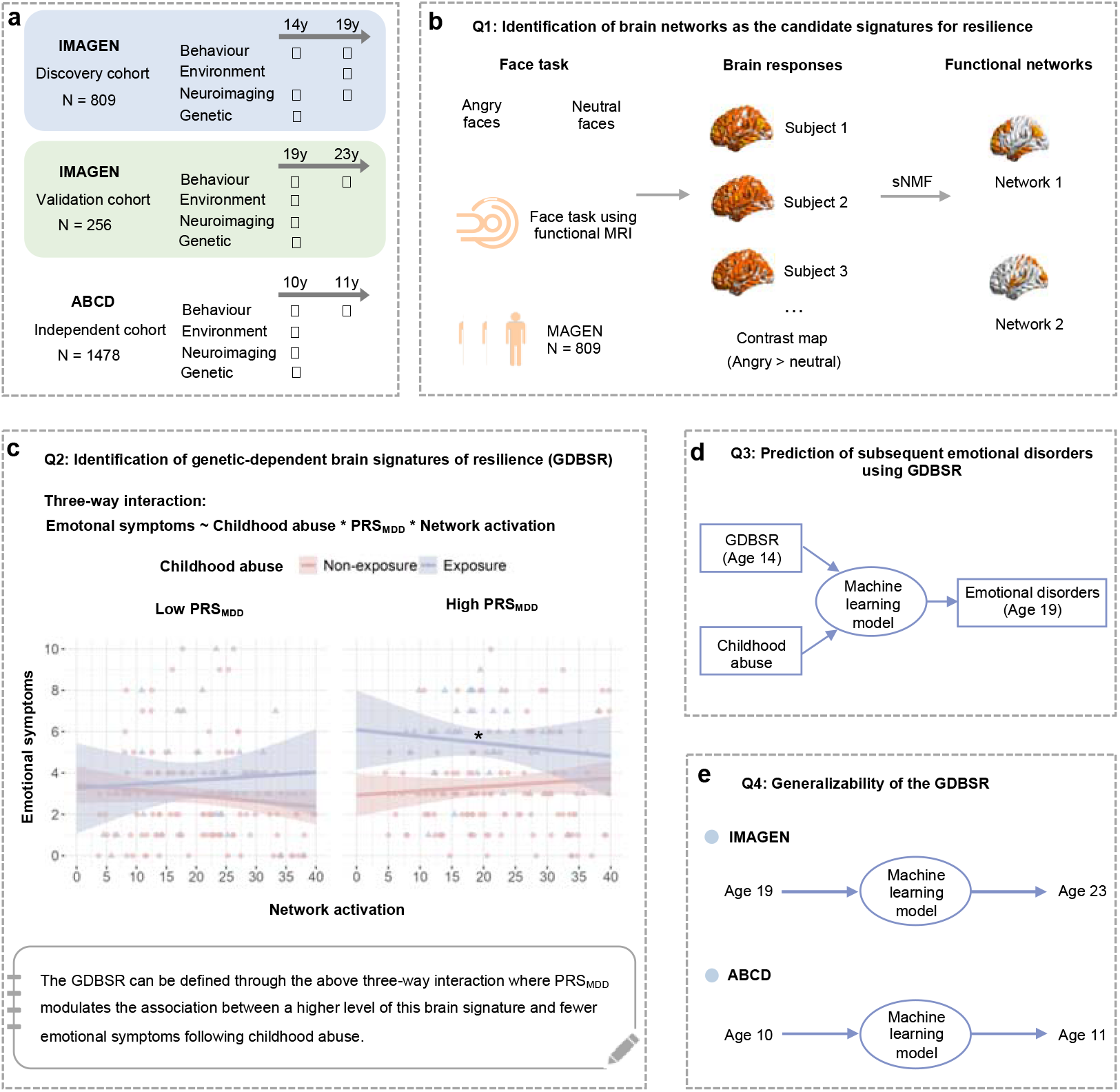
Data analysis flowchart. (a) The longitudinal cohorts used in this study. (b) We isolated distinct functional networks as candidate signatures for resilience in the IMAGEN cohort at age 19 on the basis of brain responses to angry faces using the sparse non-negative matrix factorization (sNMF). (c) We identified the genetic-dependent brain signature of resilience (GDBSR) by detecting the three-way interaction among the candidate networks in (b), childhood abuse and PRS_MDD_ in relation to emotional symptoms. (d) We tested the predictability of the GDBSR using machine learning models. (e) We checked the generalizability of the GDBSR to another developmental stage and an independent dataset.

## Results

### Summary of experimental steps

Using a large longitudinal sample of adolescents at ages 14.42±0.41 and 19.02±0.75 years old (i.e., the IMAGEN cohort^18^, N=809, 430 girls), we first decomposed brain responses to angry faces into distinct functional networks as the candidate signatures for resilience by sparse non-negative matrix factorization (sNMF). We also characterized these networks in terms of neuroanatomy, function, development, and sex difference. Second, we examined the genetic modulation of the resilience-brain associations by testing the three-way interaction on emotional symptoms, involving the candidate networks, childhood abuse and polygenic risk score for depression (PRS_MDD_). The GDB signatures of resilience can be identified when the PRS_MDD_-by-network reduces the impact of childhood abuse on emotional symptoms. Third, we built prediction models using the identified GDB signature of resilience at age 14 to predict emotional disorders at age 19. Finally, we tested the generalizability of the prediction models using both the latest follow-up data at age 23 in the IMAGEN cohort and another independent cohort, namely the Adolescent Brain Cognitive Development (ABCD) cohort ^20^ (Figure 1).

### Identification of two functional networks as candidate signatures for resilience

The brain’s emotion processing system was activated by an fMRI face task^18^. We analyzed the angry>neutral contrast map for activations responding to angry faces higher than those to neutral faces (Figure 1b). By applying the sNMF with optimal parameters to these brain activation data (Figure S2), we identified two distinct networks as the candidate signatures for resilience, including the orbitofrontal- and occipital-related networks (Figures 2b). The orbitofrontal-related network mainly covered the lateral orbitofrontal cortex (OFC), ventromedial prefrontal cortex (vmPFC), medial superior prefrontal cortex, anterior cingulate cortex (ACC), precuneus, posterior cingulate cortex and dorsolateral prefrontal cortex (dlPFC). The occipital-related network was mainly located in visual cortical regions: the lingual gyrus, cuneus, part of the inferior occipital gyrus (including the occipital face area, OFA), fusiform gyrus (including the fusiform face area, FFA), insula, amygdala, and Heschl’s gyrus (Figure 2c, Table S1). Using a database of brain functions (i.e., the NeuroSynth), we found that the orbitofrontal-related network was mainly related to high-level cognitive terms, such as episodic memory, memory retrieval and self-reference, while the occipital-related network showed associations with perceptual terms, such as vision and perception (Figure 2d). Furthermore, by conducting gene set enrichment analysis, we found that the orbitofrontal-related network but not the occipital-related network was associated with the dopaminergic synapse pathway (Figure S4, S5).

**Figure 2.**
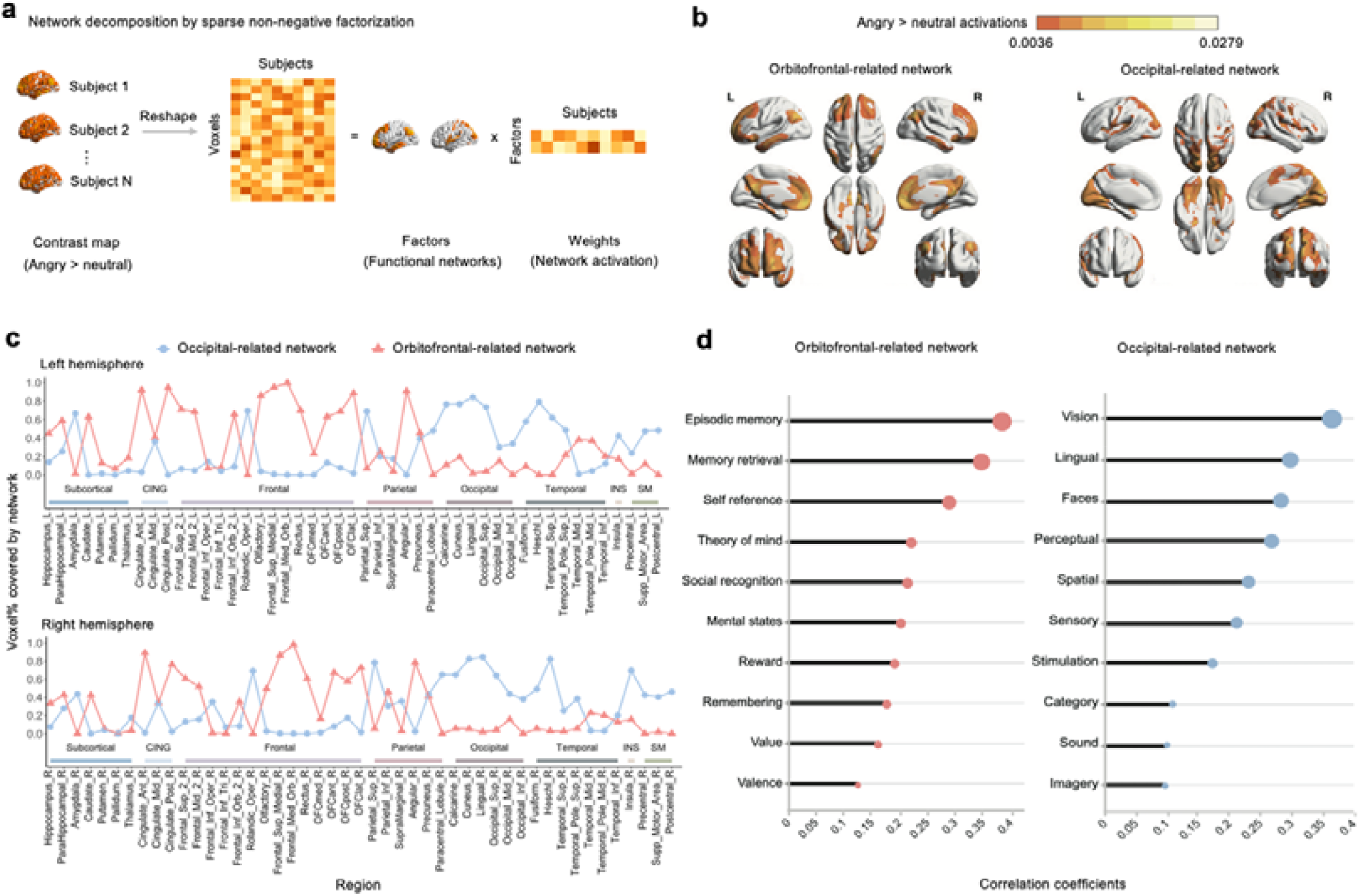
Identification of two networks as candidate signatures of resilience. (a) Brain responses to angry faces were decomposed into functional networks and corresponding network activation. (b) Brain maps represent the orbitofrontal-related network and the occipital-related network. The bright color indicates a high contribution at the spatial location of the network. (c) The voxel proportion of AAL2 regions covered by these two networks. CING, cingulate cortex; INS, insula; SM, sensorimotor. (d) NeuroSynth decoding of the networks. The lollipop charts show the correlation coefficients for each network with the top 10 terms.

### Sex differences in these networks

Sex differences in neurodevelopmental patterns of the brain’s emotion processing system may yield distinct brain signatures of resilience for boys and girls ^21^. Therefore, we explored the sex differences of the candidate signatures for resilience and found significant sex differences in these two networks at age 19 years and in their developmental trajectories between ages 14 and 19 years. Compared with boys at age 19, we found that the network activation (i.e., the factor weight) of the occipital-related network was smaller in girls (*β* = -0.230, 95% CI = [-0.369, -0.090], p = 0.001; Table S2). During the 5-year follow-up period, we found that the activation of the orbitofrontal-related network increased in both boys 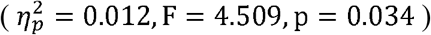 and girls 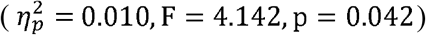. Meanwhile, the activation of the occipital-related network significantly increased in boys 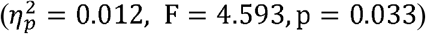 but not in girls (p = 0.643; Tables S3-4). Our results highlighted the importance of exploring the brain signatures of resilience for boys and girls respectively.

### Genetic-dependent brain signatures of resilience

As expected, higher levels of childhood abuse were associated with more emotional symptoms at age 19 in both boys (*β*=0.205, 95% CI=[0.091, 0.319], p=0.0004, N=379) and girls (*β*=0.146, 95%CI=[0.059, 0.234], p=0.001, N=430). Indeed, we found significant three-way interactions among childhood abuse, PRSMDD, and both the activations of the orbitofrontal-related (W=0.989, p=0.159 in the Wilk-Shapiro test; *β*= -0.128,95% CI = [-0.224, -0.031],p = 0.009 for the linear regression model) and the occipital-related networks (W=0.980, p=0.118 in the Wilk-Shapiro test; *β*= -0.148,95% *CI*[-0.253, -0.043], *p* = 0.005 for the linear regression model) in predicting emotional symptoms in girls at age 19 (Table S5). For illustration purposes, childhood abuse was binarized by clinical cut-offs to indicate exposure and non-exposure. High and low PRS_MDD_ were determined by a median split, as were high and low network activation. Decomposing the interaction concerning the orbitofrontal-related network revealed that among the individuals carrying high PRS_MDD_, higher activation of this network was associated with fewer emotional symptoms following childhood abuse (Figure 3a). Therefore, among girls, high PRS_MDD_ together with high activation of the orbitofrontal-related network defined a GDBSR. Similarly, we found that low PRS_MDD_ together with low activation of the occipital-related network defined another GDBSR for girls (Figure 3b). No such genetic modulations were significant in boys, and therefore we focused on girls in the following analyses.

**Figure 3.**
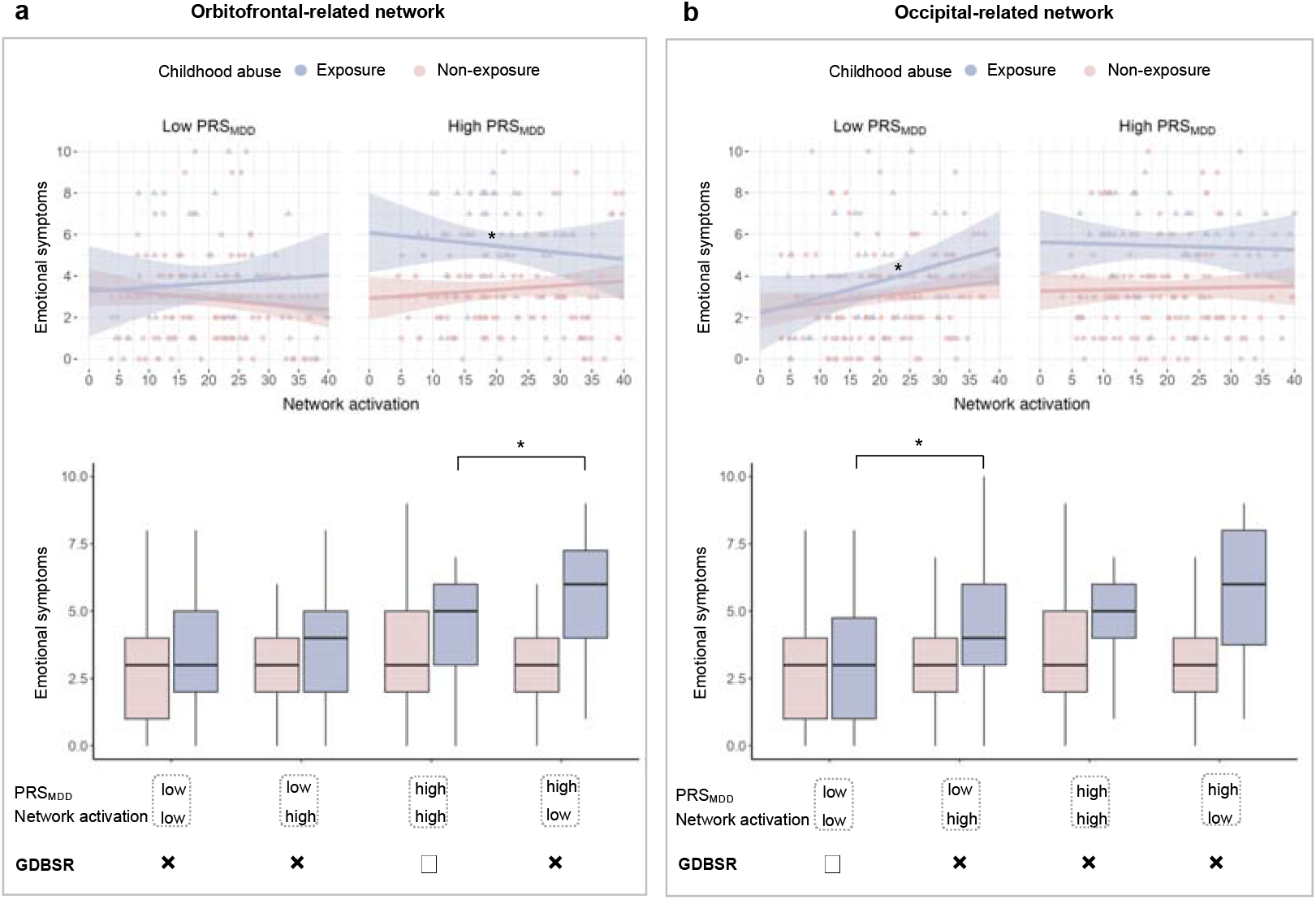
Identification of the GDBSR. Three-way interaction effects. For illustration purposes, childhood abuse was dichotomized into exposure and non-exposure based on clinical cut-offs (Methods). PRS_MDD_ levels were categorized into high and low using a median split. In the bottom of each panel, network activation was also dichotomized into high and low using a median split. (a) The GDBSR was identified by high activations of the orbitofrontal-related network together with high PRS_MDD_. (b) The GDBSR was defined by low activations of the occipital-related network together with low PRS_MDD_. * represents p<0.05.

### Sensitivity analyses

The three-way interactions identified above remained significant in the following sensitivity analyses. First, these interactions were confirmed when the childhood abuse was binarized by clinical cut-offs (Table S6). Second, these interactions remained significant after additionally controlling for the age, childhood neglect, IQ and substance use (Table S7). Next, these interactions were specific to emotional symptoms only and were not significant for the other four types of behavioral problem scores in the SDQ. Finally, these interactions on the emotional symptoms were specific to PRS_MDD_ and were not significant for either PRS_ADHD_ or PRS_SCZ_ (Table S8).

### Prospective analyses of the genetic-dependent brain signature of resilience

We used the cross-lagged panel model to delineate the directionality in the associations between network activations and emotional symptoms. After adjusting for both childhood abuse and PRS_MDD_, we found only one significant directionality in girls from the orbitofrontal-related network at age 14 to emotional symptoms at age 19 (*β*= 0.015,95% CI = [0.002, 0.027]; Figure S6). This finding was confirmed by the prospective prediction model using this network at age 14 to predict the increase in emotional symptoms during the 5-year follow-up period (*β*= 0.128,95% CI = [0.029, 0.227]; *p* = 0.010 by 1000 permutations), but not the other way around (Figure S7; Table S9). In summary, these results implied the potential predictability of the orbitofrontal-related network for emotional disorders following childhood abuse for individuals carrying high genetic risks for depression.

### Prediction of emotional disorders using the genetic-dependent brain signature of resilience

We built machine learning models (i.e., the support vector machine) using data at age 14 to predict emotional disorders at age 19 (See Methods for more details). The baseline model considered the following variables: childhood abuse, emotional symptom score, sites of data collection, handedness, pubertal status, socioeconomic status, and BMI. Based on the orbitofrontal-related GDBSR identified above, we also built a GDBSR model by adding the network activation and its interaction with childhood abuse into the baseline model. By the 5-fold cross-validation with 10 repetitions, we found that among girls with high PRS_MDD_ (*N* = 215, of whom 105 were cases) using the GDBSR model outperformed the baseline model (*AUC*:0.757 ± 0.059, *ΔAUC* = 0.016; *t*_49_= 3.462, *p* = 0.001;Table 2). As a control condition, the GDBSR model could not improve the prediction accuracy for the girls with low polygenic risks for depression (*N* = 215, of whom 85 were cases; Table 2).

### Prediction model was extended to early adulthood

To test whether the predictability of the GDBSR for emotional disorders can be extended to early adulthood, we used the data at age 19 to predict emotional disorders age 23 in the IMAGEN cohort. We confirmed that the GDBSR model again outperformed the baseline model for girls with high PRS_MDD_ (N=128, of whom 63 were cases; *AUC*:0.748 ± 0.014, *ΔAUC* = 0.011; *t*_49_= 8.563, *p* < 0.001; Table 2).

### Prediction model was generalizable to the ABCD cohort

To test the generalizability of the above finding to independent samples, we used the population-based ABCD cohort^20^. Applying the matrix factorization established above using the IMAGEN sample to the brain activations measured by the negative>neutral contrast of the EN-back task in the ABCD cohort^22^, we estimated the activations of the orbitofrontal- and occipital-related networks. Again, as compared with the baseline model, the GDBSR model using the orbitofrontal-related network at age 10 improved the prediction of emotional disorders at age 11 among the girls with high polygenic risks for depression (N=739, of whom 118 were cases; *AUC*:0.856 ± 0.035, *Δ AUC* = 0.009; *t*_49_= 4.248, *p* < 0.001; Table 2).

## Discussion

Using a discovery sample, a validation sample and another independent test cohort, the current study revealed genetic-dependent brain signatures of resilience. To identify functional networks as candidates for brain signatures of resilience, we used the sNMF approach and decomposed the brain responses to angry faces into the activations of only two distinct networks, the orbitofrontal- and occipital-related networks. These networks had different developmental patterns and significant sex differences. For girls, but not boys, we found two GDBSR, including one defined by high activations of the orbitofrontal-related network together with high polygenic burden for depression, and the other one defined by low activations of the occipital-related network together with low polygenic burden for depression. We found only the orbitofrontal-related signature had the prospective association with emotional symptoms, and this signature at age 14 predicted emotional disorders at age 19. Notably, this prediction was extendable into early adulthood and generalizable to another independent cohort. These findings highlighted the genetic modulation of the orbitofrontal function for resilience, laying the foundation for enhancing adolescent mental health through resilience promotion.

Our findings discovered two separable and interacting networks processing the angry facial expressions in adolescents. Existing literature has hypothesized that there are multiple interconnected emotional circuits in the brain for facial emotion processing^23^, and these systems have hierarchically developmental trajectories during adolescence^24^. Here, combined a longitudinally functional neuroimaging sample of the emotional face task for adolescents with an advanced matrix factorization approach, we identified a two-network system underlying the angry face processing. Many key parts of the orbitofrontal-related network, including the vmPFC^25^, the ACC^26^ and the lateral OFC^27^, have long been implicated in the neural representations of negative emotion^28^. Notably, this network covering more than 80% of the lateral OFC but less than 23% of the medial OFC (Table S1) provided a strong evidence supporting the theory of the positive-to-negative gradient in the medial-to-lateral OFC^29^. Meanwhile, the occipital-related network is well supported by a 2022 meta-analysis of 141 fMRI studies showing the occipital cortex as a key part of the facial emotion processing system^30^. Longitudinally, the medial prefrontal activity in the orbitofrontal-related network implicated in emotion regulation grows throughout adolescence^31^, while the occipital activity including those in the face-selective regions (i.e., the fusiform gyrus) in the occipital-related network often shows substantial developmental changes before adolescence^32^. These changes in the two-network facial emotion processing system may confer some adaptive advantages, such as greater flexibility in adjusting one’s intrinsic motivations and goal priorities amidst changing social contexts in adolescence.

The current findings emphasize the key role of genetic modulations in the brain’s capability of resilience. Previous studies have reported inconsistent findings on the relationship between the brain’s facial emotion processing system and resilience ^10,11^. This inconsistency may be partially explained by our finding of the genetic modulation. Such modulation is not so surprising as the genetic risks for depression have already been associated with both structures and functions of the brain’s facial emotion processing system^33^. Our finding of the resilience-related advanced maturation of the orbitofrontal function provided strong evidence of the stress acceleration hypothesis for resilience^34^. The stronger function of the orbitofrontal-related network, including the dlPFC, OFC and hippocampus, may be linked to resilience through a better neurocognitive function of the top-down suppression of traumatic memories^35^. This link was further supported by a clinical rTMS study of patients with MDD, where depression symptoms were ameliorated through enhanced activations in both OFC and hippocampus^36^. This is also supported by the overlap between this network and the default mode network (DMN), particularly medial frontoparietal regions, which have been implicated in remembering the past and self-referencing ^37^. In an imaging genetic study, the alterations of the DMN have been associated with both childhood trauma and the gene expression of SLC6A4^38^. Furthermore, our enrichment finding of the dopaminergic synapse pathway provided a neurobiological link between the orbitofrontal-related network and the dopaminergic signature of resilience^39^. Our finding of non-significant three-way interactions in boys may be due to the fact that boys have fewer emotional symptoms at age 19 when compared with girls (*β*=-0.668, 95%CI=[-0.798, -0.537], p<0.001 in the IMAGEN sample)^40^.

Our findings also have significant clinical implications for promoting adolescent mental health. One step beyond the association, the unidirectional cross-lagged association from the orbitofrontal-related network to emotional symptoms indicated the possibility of building resilience through enhancing the function of this network. Our findings using the validation sample and the independent sample further show that the time window for this intervention is open at least from preadolescence to late adolescence. Recently, neurofeedback trainings, such as the real-time fMRI feedback training of OFC^41^ and amygdala^42^, have been used to enhance emotion regulation skills and reduce emotional symptoms. However, the intervention results are mixed. Our findings suggest that the OFC-targeted interventions might be particularly effective for those individuals carrying high genetic risks for depression. Therefore, the genetic-informed and neuroimaging-targeted approach might offer a promising way of promoting adolescent mental health.

The current study is not without limitations. First, we focused only on the brain function of the facial emotion processing. Future studies are needed to test the generalizability of our findings to other types of emotional processing, which might lead to the discovery of additional brain signatures for resilience. Second, apart from the covariates considered in the current study, many other psychosocial and environmental factors (e.g., intervention program, school engagement, etc.) can also contribute to the recovery from the exposure to childhood abuse^43^. Future researches with comprehensively characterized information of these factors are needed to assess the effects of these factors on resilience. Third, the clinical value of building resilience through the genetic-informed and neuroimaging-targeted intervention strategy needs to be confirmed by randomized clinical trials.

Taken together, our study uncovered genetic-dependent brain signatures of resilience. This work emphasizes that the brain mechanisms underlying resilience might be better understood in the context of environment-gene-brain interactions.

## STAR Methods

### Participants

Participants were drawn from the IMAGEN project, a multicenter longitudinal study of adolescent brain development and mental health that recruited 2000 participants in Europe and the UK^18^. This study involves the data of each participant at ages 14 and 19. After quality control, 809 adolescents (430 girls) with complete neuroimaging data and behavioral scores at both ages 14.42±0.41 and 19.02±0.75 years old were included in this study (Table 1; Figure S1). The local research ethics committees approved this study, and written consent was obtained from each participant and a parent or guardian.

**Table 1.** Demographic characteristics of the IMAGEN sample in this study.

| Variables | Age 14 (N=809) | Age 19 (N=809) |
| --- | --- | --- |
| Age, Years | 14.42 ($\pm 0.41$) | 19.02 ($\pm 0.75$) |
| Sex, Male, n (%) | 379 (46.85%) | 379 (46.85%) |
| BMI | 20.59 ($\pm 3.19$) | 22.76 ($\pm 3.97$) |
| Social Economic Status | 2.61 ($\pm 2.39$) | 2.61 ($\pm 2.39$) |
| Hand, Right, n (%) | 696 (86.03%) | 696 (86.03%) |
| Research Site (%) |  |  |
| London | 146 (18.05%) | 146 (18.05%) |
| Nottingham | 107 (13.23%) | 107 (13.23%) |
| Dublin | 51 (6.30%) | 51 (6.30%) |
| Berlin | 73 (9.02%) | 73 (9.02%) |
| Hamburg | 111 (13.72%) | 111 (13.72%) |
| Mannheim | 97 (11.99%) | 97 (11.99%) |
| Paris | 117 (14.46%) | 117 (14.46%) |
| Dresden | 107 (13.23%) | 107 (13.23%) |
| Pubertal status | 4.14 ($\pm 0.97$) | |
| Childhood abuse | 2.49 ($\pm 4.04$) | 2.49 ($\pm 4.04$) |
| PRS<sub>MDD</sub> | -0.001($\pm 8e-05$) | -0.001($\pm 8e-05$) |
| Emotional symptoms | 2.63 ($\pm 2.05$) | 2.82 ($\pm 2.31$) |

BMI, body mass index; PRS<sub>MDD</sub>, Polygenic risk scores for major depression disorder.
Numbers of subjects are presented as integers (percentage), and quantitative
measurements are presented as mean values ± standard deviations.

**Table 2.**
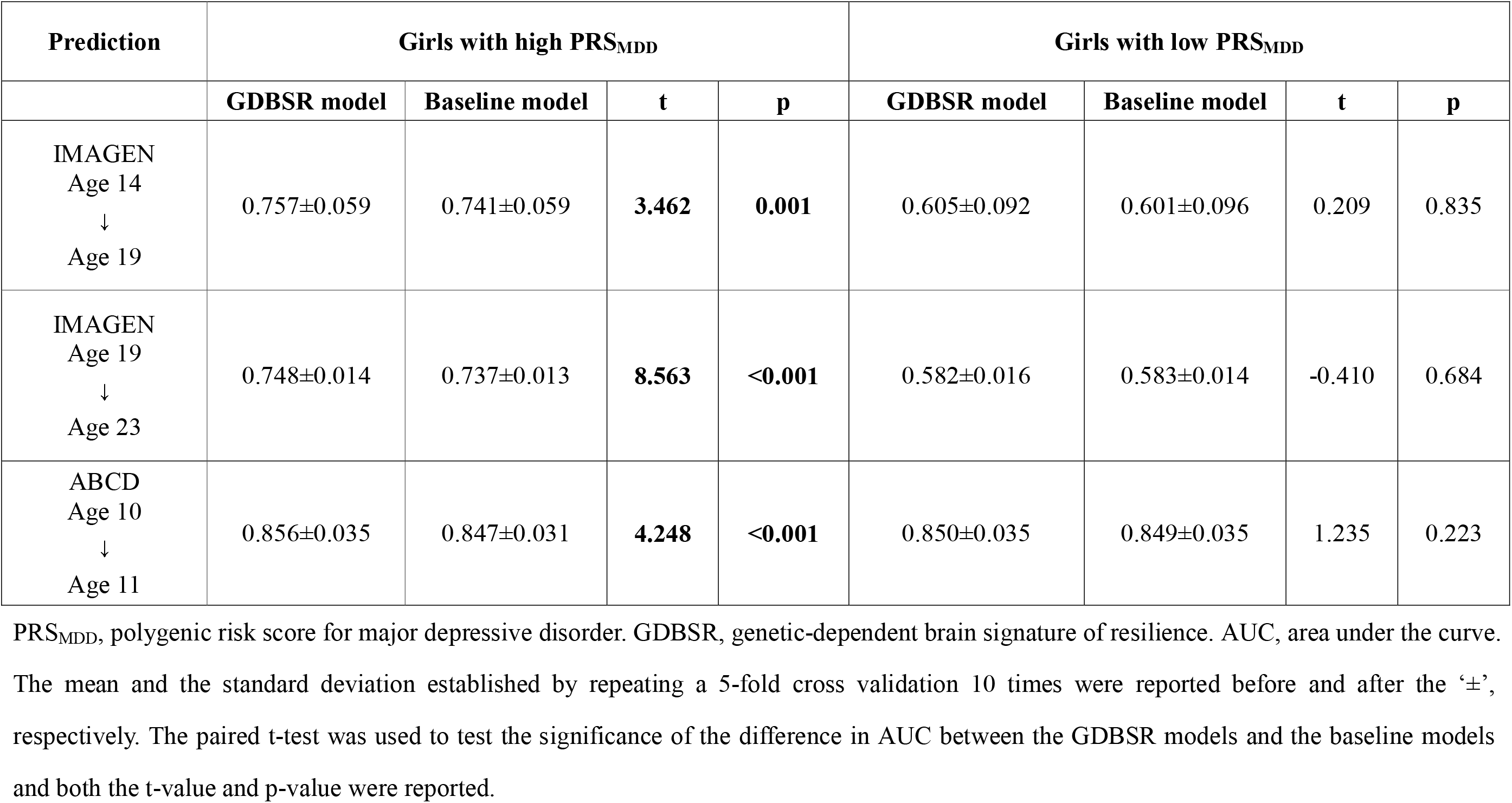
Comparison of model performance for the prediction of emotional disorders in girls.

### Measurements

#### Behavioral and emotional problems

The Strengths and Difficulties Questionnaire (SDQ) is a valid and reliable assessment and is often used to measure the emotional and behavioral problems in adolescents, including emotional symptoms, conduct problems, hyperactivity/inattention, peer relationship problems, and prosocial behavior^44^. SDQ questionnaires gathered directly from adolescents themselves are more reliable than those from their parents, especially for the emotional symptom subscale ^45^. Therefore, the self-reported versions of the SDQ at ages 14 and 19 were used in this study.

#### Childhood abuse measurements

The Childhood Trauma Questionnaire (CTQ^46^) is a 28-item self-report inventory used to assess the history of abuse and neglect before the age of 19 years. Since the IMAGEN study focused on a population-based cohort, the severity of each type of abuse may be underestimated. Therefore, three abuse subscales (*i*.*e*., emotional abuse, physical abuse and sexual abuse) were summed to generate a composite measure of childhood abuse^47^. The higher the abuse score, the greater the severity of childhood abuse.

#### Polygenic risk scores

Since emotional disorders are not single-gene diseases, it is promising to use PRS to reflect the complex genetic architecture in the context of environment-gene-brain interactions ^7^. We used the GWAS summary data provided by the Psychiatric Genomics Consortium as the discovery sample. 493,592 single nucleotide polymorphisms (SNPs) were shared by the discovery sample and the IMAGEN cohort. After the quality control measures (Method S1), a total of 123,481 SNPs were selected to compute the PRS_MDD_ in our sample using the genetic analysis tool PLINK. The means of the PRSs at 7 p-value thresholds (*i*.*e*., 0.001, 0.05, 0.10, 0.20, 0.30, 0.40, and 0.50) were used in the current study in keeping with a previous study ^48^.

#### Nuisance covariates

Pubertal status was assessed using the Pubertal Development Scale. A total neglect score was generated from the summation of two types of neglect (*i*.*e*., emotional neglect and physical neglect) in the CTQ. Socioeconomic status was rated according to the total score of the family stress subsection of the Development and Well-being Assessment. The IQ score of each participant was calculated as the total score derived from the Wechsler Intelligence Scale for Children-Fourth Edition (WISC-IV). Substance use was measured using the European School Survey Project on Alcohol and Drugs (ESPAD) as ever/never smoking cigarettes, drinking alcohol, or using illicit drugs.

### The face task and fMRI preprocessing

The face task paradigm was used to elicit strong activation in the facial emotion processing system. In this task, participants passively watched 18-second blocks of either a face movie (presenting faces with angry, happy or neutral expressions) or a control stimulus (concentric circles). Details can be found in the initial report on this paradigm^19^. In this study, we explored the neural reactivity associated with angry expressions, as neuroimaging data on these expressions was available at both ages 14 and 19. After the fMRI pre-processing (Method S2), the contrast map of angry vs. neutral faces was obtained for each participant. The angry>neutral (*i*.*e*., the activations responding to angry faces were higher than those to neutral faces) activations were used to measure the activation of the facial emotion processing system in the brain responding to angry faces. Although the mechanisms underlying the neutral>angry activations remained unclear, we still examined such activations in the supplementary materials to enhance the comprehensiveness of our study. The voxels within the automated anatomical labeling (AAL2) template^49^ for grey matter were considered in the following analyses (47,640 voxels).

### Matrix decomposition

We constructed an activation matrix for the angry>neutral activations. The activation matrix has a number of rows equal to the voxel count (*m*=47,640) and a number of columns corresponding to the number of subjects (*n=*809). Sparse non-negative matrix factorization (sNMF) was employed to decompose the activation matrix at age 19 into a factor matrix and a weight matrix (Figure 2a). To facilitate meaningful sparse representation, we explicitly incorporated ℓ^0^-sparseness constraints^50^ on the columns of the factor matrix. Meanwhile, each row of the factor matrix can have only one non-zero value to ensure that no overlapping voxels among the latent factors are obtained by the decomposition (Method S3). To determine the optimal parameter for sparsity (*γ* = *L*/*m, L* is the maximal number of non-zeros voxels in each factor, *m* is the total number of voxels) and the optimal number of factors (K), we tested both the reconstruction error and the reproducibility of the obtained decompositions by a random half split for 80 times (Method S4).

### Characterization analysis of the functional networks

#### Neuroanatomical characterization

We identified the respective positions of the non-zero values in each column of the factor matrix (i.e., each latent factor) within 47,640 voxels in the AAL2 template.

#### Functional characterization

As recommended by the previous work ^51^, we compared the spatial pattern of the networks (i.e., factors) to the functional anatomy of the human brain using NeuroSynth (http://www.neurosynth.org/)^52^, an online platform for meta-analysis of functional neuroimaging literature. Specifically, we sorted all correlation coefficients for each network in descending order and adopted the top ten terms to characterize each network. Similar terms (e.g., “percept” and “perception”) were merged into a base form to avoid selecting repetitive terms.

#### Gene set enrichment analyses

To examine the neurobiological links between the identified networks and the dopaminergic signature of resilience reported in the literature^39^, we used the transcriptomic data from six neurotypical adult brains in the Allen Human Brain Atlas (AHBA) (http://human.brain-map.org)^53^. Following a preprocessing pipeline recommended by previous work (Method S5) ^54^, we obtained a 1531 (number of tissue samples from the cerebral cortex) × 15,408 (number of genes) matrix. Genes were considered significant if their expression levels differed between tissue samples inside and outside the functional networks, with a significance threshold of p <3.25 × 10^−6^ (0.05/15408). Next, we used the R packages “BiocManager” and “clusterProfiler” to identify sets of genes associated with Gene Ontology terms of biological processes and Kyoto Encyclopedia of Genes and Genomes pathway. Gene sets were considered significantly enriched with FDR q values < 0.05.

#### Sex difference

We built a linear regression model between the activation of each network (i.e., the weights of each factor) at age 19 and sex. Research sites, socioeconomic status, BMI at age 19 ^55^ and handedness^48^ were regressed out as basic covariates in this analysis and the following analyses.

#### Developmental trajectory

We applied the NMF back-reconstruction algorithm to compute the activation of each network of each participant at age 14 (Method S6). Next, for boys and girls separately, we carried out repeated measures analyses of variance (ANOVAs) to investigate the developmental trajectories of the network activations. The age 14 and age 19 network activations were the within-subject variables. In addition to the basic covariates, we incorporated pubertal status as an additional covariate, considering the relationship between pubertal maturation and the reactivity of facial emotion processing systems during early adolescence^56^.

### Modulation analysis

For boys and girls separately, associations were assessed by a linear regression model between emotional symptoms at age 19 and childhood abuse before age 19. Next, to identify the GDBSR, we examined the three-way interaction among PRS_MDD_, the activations of the above identified functional networks, and childhood abuse, in relation to emotional symptoms at age 19. The coefficient (standardized /3) of the linear regression models and its 95% confidence interval (CI) are reported. The applicability of linear model in this case was confirmed by the Shapiro-Wilk normality test for model residuals ^57^. A significant three-way interaction indicates that PRS_MDD_ modulates the association between a higher level of this brain signature and fewer emotional symptoms following childhood abuse.

### Sensitivity analyses

We tested whether the three-way interaction remained significant when the childhood abuse score was binarized using the following cut-offs as recommended in the literature^58^, including a cut-off of 8 for emotional abuse, 7 for physical abuse, and 5 for sexual abuse. If any type of the above abuse occurred, childhood exposure to abuse was scored as “1”; if not, a score of “0” was recorded. We also included age, childhood neglect, IQ or substance use as an additional covariate in the modulation models to examine their potential confounding effects. To investigate the specificity of the modulation effects, we reran the models while 1) replacing the emotional symptom scores with behavioral problem scores from the other four dimensions in the SDQ; 2) replacing the PRS_MDD_ with the PRS_ADHD_ or the PRS_SCZ_.

### Prediction models

#### Prospective associations

For significant modulation effects, we employed a two-wave cross-lagged panel model (CLPM) using the network activations and emotional symptoms at ages 14 and 19 years. In addition to the basic covariates, we incorporated BMI at age 14, pubertal status, childhood abuse and PRS_MDD_ as additional covariates, considering the potential association between emotional symptoms and both childhood abuse and PRS_MDD_. We established the 95% CI of the statistics by 1000 bootstraps. We also used linear regression models to verify such directionality (Method S7).

#### Building prediction models for late-adolescence emotional disorders

Using the networks that have significant prospective associations with subsequent emotional symptoms, we built prediction models for emotional disorders at age 19. The emotional disorders were indicated by an emotional symptom score above a clinical cut-off of 4, which has been recommended to favor the instrument’s (*i*.*e*., SDQ) sensitivity in identifying depression and generalized anxiety^59^. The high-risk group was identified as participants with above-median genetic risk for depression (*i*.*e*., PRS_MDD_>median PRS_MDD_); otherwise, the low-risk group was defined. We built the following prediction models for each group. The baseline model was a support vector machine with a linear kernel using the measurements at age 14 years, including childhood abuse, emotional symptom score, sites of data collection, handedness, pubertal status, socioeconomic status, and BMI. Next, based on the GDBSR identified above, we built the GDBSR models by adding the network activation and its interaction with childhood abuse into the baseline model. To evaluate model performance, we repeated a 5-fold cross-validation 10 times to obtain the mean area under the curve (AUC). The paired t-test was used to test the significance of the difference in AUC between the GDBSR models and the baseline models.

### Generalizability of the prediction models

#### Generalizability in early adulthood

Using the latest follow-up data at age 23 in the IMAGEN study, we tested the model performance among 256 girls. We applied the aforementioned trained models, without retraining (i.e., fixed weights), to see whether emotional disorders at age 23 can be predicted by the model using measurements at age 19.

#### Generalizability in an independent dataset

To test whether the GDBSR models could be generalized to an independent dataset, we used the data from the ABCD cohort (the ABCD data used in this study came from Data Release 5.0, http://dx.doi.org/10.15154/8873-zj65) to rerun the prediction models. This independent dataset recruited 11,875 children between 9 and 10 years of age from 21 sites across the United States^20^. The negative>neutral activations during 0 back in the EN-back task^22^ were used. We applied the NMF back-reconstruction algorithm again to compute the activations of the functional networks for each participant in the ABCD cohort. After quality control (the same as the IMAGEN cohort), 1478 participants with complete neuroimaging data, PRS_MDD_, adverse childhood experiences (ACEs)^60^, and the basic covariates at baseline, as well as the internalizing symptoms of the Child Behavior Checklist ^61^ at both baseline and the 1-year follow-up were analyzed. The emotional disorders were indicated by an internalizing symptom t score above a cut-off of 60^62^. Similarly, we first built the baseline model using the baseline measurements to predict emotional disorders at the 1-year follow-up for both the high and low genetic risk groups. Next, we added the network activation and its interaction with ACEs into the baseline model to form the GDBSR model.

## Data availability

The IMAGEN data are available by application to the consortium coordinator Dr. Schumann (http://imagen-europe.com) after evaluation according to an established procedure. The ABCD data are publicly released on an annual basis through the National Institute of Mental Health (NIMH) data archive (NDA, https://nda.nih.gov/abcd). The ABCD study data are openly available to qualified researchers for free. Access can be requested at https://nda.nih.gov/abcd/request-access. An NDA study has been created for the data used in this report under the doi: 10.15154/agv5-7v56.

## Code availability

The code used by the current study is made available at the following webpage: https://github.com/hanluyt/modulation_emotionalBrain.

## Acknowledgements

This study was partially supported by grants from the National Key Research and Development Program of China (No. 2023YFE0109700), the National Natural Science Foundation of China (No. 82272079), the Program of Shanghai Academic Research Leader (No. 23XD1423400), and the Shanghai Municipal Science and Technology Major Project (No.s: 2018SHZDZX01 and 2021SHZDZX0103). This work also received support from the following sources: the European Union-funded FP6 Integrated Project IMAGEN (Reinforcement-related behaviour in normal brain function and psychopathology) (LSHM-CT-2007-037286), the Horizon 2020 funded ERC Advanced Grant ‘STRATIFY’ (Brain network based stratification of reinforcement-related disorders) (695313), Human Brain Project (HBP SGA 2, 785907, and HBP SGA 3, 945539), the Medical Research Council Grant ‘c-VEDA’ (Consortium on Vulnerability to Externalizing Disorders and Addictions) (MR/N000390/1), the National Institute of Health (NIH) (R01DA049238, A decentralized macro and micro gene-by-environment interaction analysis of substance use behavior and its brain biomarkers), the National Institute for Health Research (NIHR) Biomedical Research Centre at South London and Maudsley NHS Foundation Trust and King’s College London, the Bundesministeriumfür Bildung und Forschung (BMBF grants 01GS08152; 01EV0711; Forschungsnetz AERIAL 01EE1406A, 01EE1406B; Forschungsnetz IMAC-Mind 01GL1745B), the Deutsche Forschungsgemeinschaft (DFG grants SM 80/7-2, SFB 940, TRR 265, NE 1383/14-1), the Medical Research Foundation and Medical Research Council (grants MR/R00465X/1 and MR/S020306/1), the National Institutes of Health (NIH) funded ENIGMA (grants 5U54EB020403-05 and 1R56AG058854-01), NSFC grant 82150710554 and European Union funded project ‘environMENTAL’, grant no: 101057429. Further support was provided by grants from: - the ANR (ANR-12-SAMA-0004, AAPG2019 - GeBra), the Eranet Neuron (AF12-NEUR0008-01 - WM2NA; and ANR-18-NEUR00002-01-ADORe), the Fondation de France (00081242), the Fondation pour la Recherche Médicale (DPA20140629802), the Mission Interministérielle de Lutte-contre-les-Drogues-et-les-Conduites-Addictives (MILDECA), the Assistance-Publique-Hôpitaux-de-Paris and INSERM (interface grant), Paris Sud University IDEX 2012, the Fondation de l’Avenir (grant AP-RM-17-013), the Fédération pour la Recherche sur le Cerveau; the National Institutes of Health, Science Foundation Ireland (16/ERCD/3797), U.S.A. (Axon, Testosterone and Mental Health during Adolescence; RO1 MH085772-01A1) and by NIH Consortium grant U54 EB020403, supported by a cross-NIH alliance that funds Big Data to Knowledge Centres of Excellence. The ABCD Study is supported by the National Institutes of Health and additional federal partners under award numbers U01DA041022, U01DA041028, U01DA041048, U01DA041089, U01DA041106, U01DA041117, U01DA041120, U01DA041134, U01DA041148, U01DA041156, U01DA041174, U24DA041123, and U24DA041147. A full list of supporters is available at https://abcdstudy.org/federal-partners.html. A listing of participating sites and a complete listing of the study investigators can be found at https://abcdstudy.org/consortium_members/. ABCD consortium investigators designed and implemented the study and/or provided data but did not necessarily participate in the analysis or writing of this report. This manuscript reflects the views of the authors and may not reflect the opinions or views of the NIH or ABCD consortium investigators. The ABCD data repository grows and changes over time. The ABCD data used in this report came from the Data Release 5.0 (http://dx.doi.org/10.15154/8873-zj65).

## Author contributions

Q.L. has full access to all the data in the study and takes responsibility for the integrity of the data and the accuracy of the data analysis. H.L. and HJ.L. designed and implemented the experiments. H.L., E.T.R. and Q.L. wrote the manuscript. T.J., C.X., S.X. preprocessed the fMRI data. Q.L., G.S. and J.F. secured funding. D.J.S., B.J.S., R.E. and N.W. provided expertise and feedback. T.B., A.B., S.D., H.F., A.G., H.G., A.H., R.B., J.M., M.M., E.A., F.N., D.O., H.L., T.P., L.P., S.H., N.H., J.H.F., M.N.S., N.V., H.W., R.W., G.W. and J.F. provided administrative, technical, or material support.

