## Supplement for "Genetic-Dependent Brain Signatures of Resilience: Interactions among Childhood Abuse, Genetic Risks and Brain Function"

Han Lu, *et al.*

### Supplementary Methods

Method S1 Genome-wide genotype data and polygenic risk scores

Whole-blood samples (~10 mL) were collected in 2,087 IMAGEN participants. The details of genotyping and quality controlling procedures can be found in a previous study(1). DNA purification and genotyping were performed by the Centre National de Génotypage in Paris. A total of 705 and 1,382 individuals were genotyped with the Illumina (Little Chesterford, UK) Human610-Quad Beadchip (582,982 SNPs) and Illumina Human660-Quad Beadchip (557,124 SNPs), both of which were based on the same Illumina HumanHap550 Genotyping BeadChip with varied additional probes, and therefore they have most of their SNPs (551,141 SNPs) in common. Genotyped calls were aligned based on assembly hg18.

To generate PRS_MDD,_ PRS_ADHD_ and PRS_SCZ_ in IMAGEN, we used GWAS summary data from the Psychiatric Genomics Consortium, which served as the discovery sample. 493,592 SNPs were shared by the discovery sample and IMAGEN. The quality control was performed by PLINK with the following steps: 1) SNPs with minor allele frequency (MAF) below 1% or imputation information score (INFO) below 0.8 were excluded from the PRS analyses; 2) SNPs with call rates ≥ 0.95 and MAF ≥ 1% were retained; 3) Individuals with high missingness rates (> 5%) and autosomal heterozygosity deviations (FHET) not within ± 3 SD were removed; 4) SNPs were filtered for call rates ≥ 0.98 and Hardy-Weinberg p values > 1e-6 (founders only) after sample quality control. Sex checks were performed to remove mismatches between biological and reported sex for subsequent analyses.

After the quality control measures, we obtained a total of 123,481 SNPs for MDD and ADHD, 89,722 SNPs for SCZ in 1843 individuals. Dosage data were converted to hard-called genotypes, and only SNPs with imputation r^2^ scores ≥ 0.3 were transferred to PRS. The genetic analyses tool PLINK was used to calculate PRSs under p-value thresholds at 0.001, 0.05, 0.10, 0.20, 0.30, 0.40 and 0.50. Then we computed the mean PRSs to obtain PRS_MDD,_ PRS_ADHD_ and PRS_SCZ_ respectively. The principal component analysis of ancestral information was performed and the first 8 principal components (PCs) were projected from the founders to other relatives. In this study, we included the first 8 PCs as covariates when PRS was considered as the predictor in the models.

Method S2 fMRI measurement and processing

Structural and functional MRI were performed on 3T MRI scanners from a range of manufacturers (Siemens, Philips, General Electrics & Bruker). All sites used the same scanning protocol; High-resolution T1-weighted structural images were acquired for anatomical localization and registration with the functional time series. Blood oxygen-level dependent (BOLD) functional images were acquired with a gradient-echo echoplanar imaging (EPI) sequence. In this task, a hundred and sixty volumes were collected per subject, each containing 40 slices parallel to the anterior–posterior bicommissural line (2.4mm slice thickness, 1mm gap, TE=30ms, TR=2200ms).

FMRI pre-processing and single-subject analyses were performed using SPM8 (<http://www.ﬁl.ion.ucl.ac.uk/spm/>). Functional time series data were first corrected for slice-timing and then corrected for movement (realigned to the first volume) and non-linearly warped on the Montreal Neurological Institute (MNI) space, using a custom EPI template. Images were then smoothed with a 5mm full-width half maximum Gaussian filter. Single-subject analyses were performed within the framework of the general linear model (GLM), using SPM default hemodynamic response function.

After the fMRI pre-processing, the contrast map of angry vs. neutral faces was obtained for each participant (53 × 63 × 46 voxels, at a resolution of 3 × 3 × 3 mm^3^). And then each contrast map was separated by the non-negative and the negative values (converted to positive values) for the angry > neutral (*i.e.*, the activations responding to angry faces were higher than those to neutral faces) and neutral > angry activations, respectively. For each type of activation, we built an activation matrix for 47640 voxels (*i.e.*, rows) and 809 participants (*i.e.*, columns).

Method S3 Non-negative matrix factorization (NMF)

NMF is an unsupervised machine learning technique used to find part-based, linear representations of non-negative data(2). It has recently emerged as a promising approach to capture reliable and robust low-dimensional representation from high-dimensional input data and has been widely used in MRI analysis(3-5). Compared with other matrix factorization methods such as principal components analysis and vector quantization, NMF can learn more intuitive and meaningful part-based representation as NMF does not allow negative entries in the matrix decomposition. Therefore, NMF is well-suited for functional MRI to decouple complex interactions among various brain networks into latent factors and to measure the contribution of each latent factor to the brain activation in each participant.

NMF aims to factorize a non-negative data matrix X into a product of non-negative matrices W and H. Here, X is the activation matrix with dimensions *m×n* (where *m* is the number of voxels and *n* is the number of subjects). W is the factor matrix with dimensions *m×k* (where *k* is the number of factors) and H is the weight matrix with dimensions *k×n*. In this study, each column of the factor matrix represents the contribution of each voxel to a latent factor of brain activation. Each row of the weight matrix represents the contribution of a latent factor to the brain activation in each participant. To facilitate the meaningful sparse representation, we explicitly incorporated $\mathcal{l}^{0}$-sparseness constraints(6) on the columns of the factor matrix. Meanwhile, each row of the factor matrix can have only one non-zero value, to ensure that no overlapping voxels among the latent factors obtained by the decomposition. We implemented NMF with Frobenium norm as the objective function. This optimization problem is defined as follows:

$$minimize \left\| \boldsymbol{X}-\boldsymbol{WH} \right\|_{F}$$

$$subject to \boldsymbol{W(:)}\geq0,$$

$$\boldsymbol{H(:)}\geq0,$$

$$\left\| \boldsymbol{w}_{i} \right\|_{0}\leq L, \forall i$$

$$\left\| \boldsymbol{W}\left( j, : \right) \right\|_{0}\leq1, \forall j$$

$$j is the row index of \boldsymbol{W}$$

where$\left\| \cdot\right\|_{F}=\sqrt{\sum_{i,j} \left| x_{i, j} \right|^{2}}$, ≥ denotes the element-wise greater-or-equal operator, sparsity parameter $L\mathbb{\in N}$ is the maximal allowed number of non-zero entries in the$\boldsymbol{w}_{i}$ (*i.e.*, the i^th^ column of **W**) and $\left\| \cdot\right\|_{0}$is the $\mathcal{l}^{0}$-norm.

We used fast combinatorial approach for nonnegative least squares(7) to obtain factors and weights, which is an efficient and powerful technique for solving the NMF problem. The package for NMF is available at <https://github.com/smatmo/l0-sparse-NMF>. NMF was first applied on the activation matrices at age 19 years old.

Method S4 Determining the optimal parameter for sparsity and the optimal number of factors

In the NMF framework with sparseness constraint, increasing sparsity improves the interpretability and specificity but influences the performance of NMF. In order to determine the sparsity parameter ($\lambda=L/m$, *L* is the maximal number of non-zeros voxels in each factor, $\lambda$ from 0.1 to 0.9) and the optimal number of factors (*K* from 2 to 9), we tested both the reconstruction error and the reproducibility of the obtained decompositions. This strategy is guided by the previous work(5).

We investigated the reconstruction error of the matrix decomposition of brain activations with the hyperparameters $\lambda$ and *K*. The reconstruction error was defined as follows:

*Reconstruction error =* $\left\| X-WH \right\|_{F}= \sqrt{\sum_{i,j} {(X_{ij}-{(WH)}_{ij})}^{2}}$

Where X is the activation matrix, W is the factor matrix and H is the weight matrix. This measurement was used to evaluate the accuracy of the matrix decomposition.

To test the reproducibility of the obtained decompositions, we randomly split the subjects at age 19 years old into two subgroups and reran the NMF model separately. Reproducibility was quantified by the similarity of factors between these two subgroups for each specific lambda and K. We computed the mean correlation between corresponding factor pairs (using the Hungarian matching algorithm).

We repeated the NMF procedure 80 times to weaken the influence of the random initializations. The optimal parameter for sparsity and the optimal number of factors were obtained by considering smaller reconstruction error and larger mean correlation simultaneously.

Method S5 Gene set enrichment analysis

To examine the neurobiological links between the factors and the dopaminergic signature of resilience reported in the literature(31), we used the transcriptomic data from six neurotypical adult brains in the Allen Human Brain Atlas (AHBA) ( <http://human.brain-map.org>)(8). In this database, gene expressions were normalized across all samples and genes using BF normalization methods. We followed the AHBA pre-processing pipeline suggested by previous work(9), including the following steps:

1) Probe-to-gene re-annotation: This reannotation was done by using the reference genome assembly GRCh38.p12 (released in 2017/12). Following the research of Shen et al2, 45,461 (77.5%) probes were annotated to unique genes. This re-annotated set of 45,461 probes corresponded to 19,951 unique genes.

2) Data filtering: The AHBA provided a binary indicator for those expressions did not exceed the background in at least 50% of all cortical and subcortical samples across all subjects. Excluding these probes gave a set of 31,342 probes and 15,409 unique genes.

3) Probe selection: As recommended by Arnatkevic̆iūtė et al, expressions were averaged among probes assigned to the same gene. As reported in our previous publication2, the mean approach and the max approach gave highly correlated gene expressions (r=0.88).

4) Using the Harvard-Oxford atlas, the samples were separated into the cortical and the subcortical areas. Only the samples in the left hemisphere were used in the following analyses, since the samples in the right hemisphere were collected from only 2 of the 6 donors.

5) Normalization: The expression data were first normalized within sample and across-gene, and then normalized across samples. Given the systematic differences in gene expressions between the cortical and the subcortical areas, the normalizations were conducted separately for the samples from the subcortical and the cortical tissues. One gene failed the normalization and therefore was deleted, resulting 15,408 unique genes.

After preprocessing, we obtained 1531 tissue samples in the cortex, each with Montreal Neurological Institute (MNI) coordinates and expression values for 15,408 genes. To determine the number of tissue samples located inside and outside the factors, we defined a spherical region of interest (ROI) with a radius of 5mm centered at MNI coordinates of each gene expression sampling site. We then calculated the proportion of voxels within this ROI that fell inside the factor. If the proportion exceeded 50%, the tissue sample was considered to be inside the factor. Genes were considered significant if their expression levels differed between tissue samples inside and outside the factors using t-test, with a significance threshold of p <3.25$\times$10-^6^ (0.05/15408). Next, we used the R packages “BiocManager” and “clusterProfiler” to identify sets of genes associated with Gene Ontology (GO) terms of biological processes and Kyoto Encyclopedia of Genes and Genomes (KEGG) pathway. Gene sets were considered signiﬁcantly enriched with FDR q values < 0.05. We reported the top 10 significant GO terms and KEGG pathways, respectively in Figure S4-5.

.

Method S6 NMF back-reconstruction algorithm

After the factors were obtained by matrix decomposition of brain activations at age 19 years old, we used NMF back-reconstruction algorithm to reconstruct the activation matrices at age 14 years old. Specifically, the factor matrix at age 19 years old was used to estimate the weight matrix at 14 years old. The procedure was defined as follows:

$$X_{n>a activations at age 19}= W_{n>a activations at age 19}H_{n>a activations at age 19}$$

$$X_{n>a activations at age 14}=W_{n>a activations at age 19}H_{n>a activations at age 14}$$

$$X_{a>n activations at age 19}= W_{a>n activations at age 19}H_{a>n activations at age 19}$$

$$X_{a>n activations at age 14}=W_{a>n activations at age 19}H_{a>n activations at age 14}$$

Fast combinatorial approach for nonnegative least squares(7) was used to solve NMF problem and estimate the H matrix. The difference between NMF and NMF back-reconstruction is that the latter iteratively updates the H matrix, while keeping the W matrix obtained from matrix decompositions at age 19 years old fixed, and does not stop until the reconstruction error is almost unchanged compared with the previous iteration.

Method S7 Prospective prediction model

We used linear regression models to predict the rate of change in emotional symptoms between ages 14 and 19 as $(E_{19}-E$_14_$) / (E_{14}+1)$ by each of the factor weight (that is, network activation) at age 14 (F_14_). The following covariates were considered: rate of change in factor weight$(F_{19}-F_{14}) / (F_{14}+1)$, childhood abuse, PRS_MDD_ and their three-way interaction term, socioeconomic status, sites of data collection, pubertal status, handedness, BMI_14_ and E_14_.

In the opposite direction, we used the baseline emotional symptom score (E_14_) to predict the rate of change in the factor weight. The rate of change in the emotional symptom score, childhood abuse, PRS_MDD_ and their three-way interaction, socioeconomic status, sites of data collection, pubertal status, handedness, BMI_14_ and F_14_ were used as covariates in the models.

For the significant findings survived the FDR correction, the permutation test (n=1000, p_PERM_<0.05) was used to confirm this significance, to exclude the potential influence of the non-normality of the data distribution on the significance given by the linear model.

### Supplementary Figures

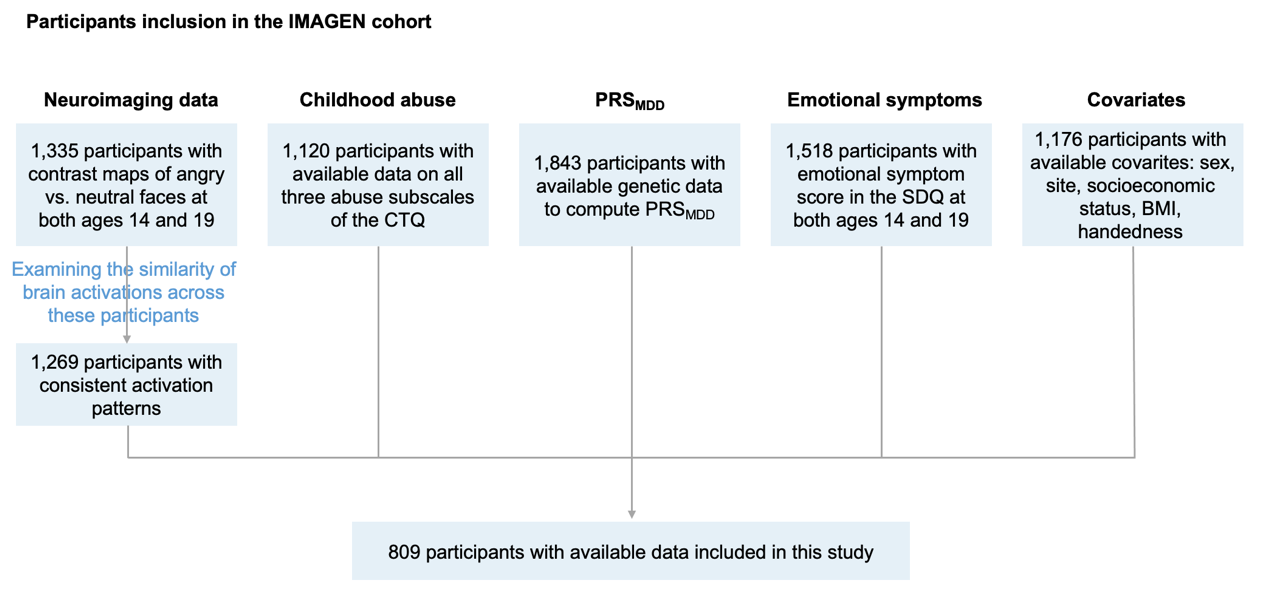

Figure S1 Flow diagram of the participants in the IMAGEN cohort included in this study

The Childhood Trauma Questionnaire (CTQ) was available at age 19 but not at age 14.

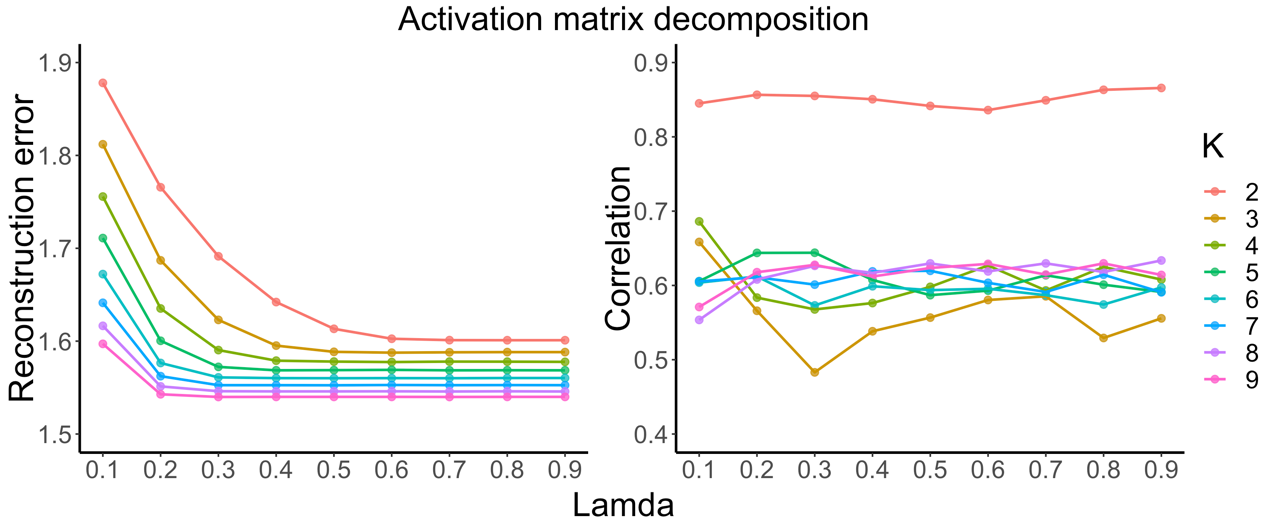

Figure S2 Reconstruction error and reproducibility for matrix decomposition of angry>neutral activations.

The sparsity parameter $\lambda$=L/m, L is the maximal number of non-zeros voxels in each identified network, m is the total number of voxels, $\lambda$ from 0.1 to 0.9; The number of factors K from 2 to 9. According to the optimal criterion for matrix decomposition, we choose $\lambda=0.3, K=2$for this activation. Therefore, we identified **two** latent factors (that is, networks) for the angry>neutral activations.

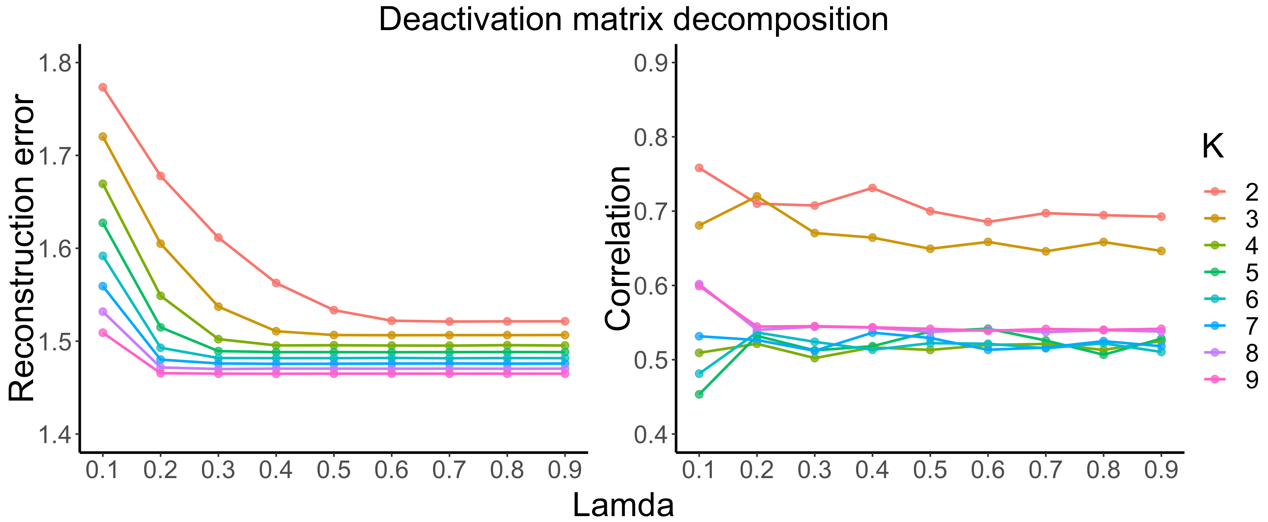

Figure S3 Reconstruction error and reproducibility for matrix decomposition of neutral>angry activations

The sparsity parameter $\lambda$ =L/m, L is the maximal number of non-zeros voxels in each identified network, m is the total number of voxels, $\lambda$ from 0.1 to 0.9; The number of factors K from 2 to 9. According to the optimal criterion for matrix decomposition, we choose $\lambda=0.2, K=3$for this activation. Therefore, we identified **three** latent factors (that is, networks) for the neutral>angry activations.

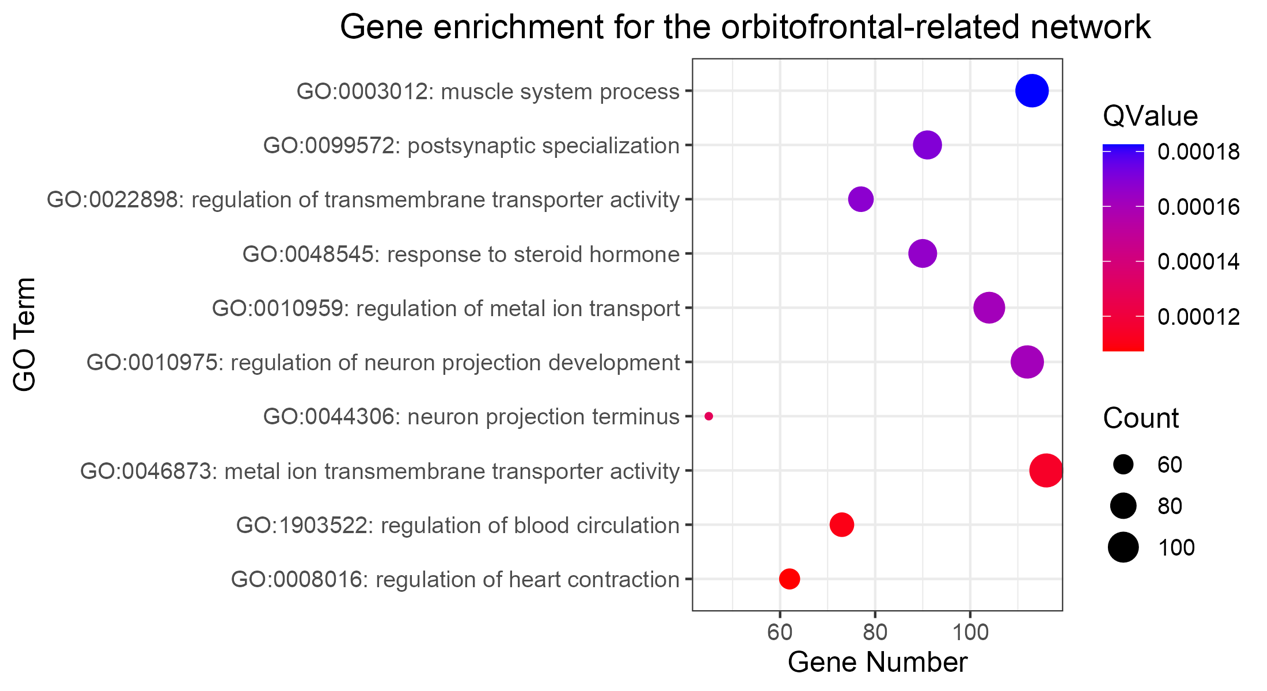

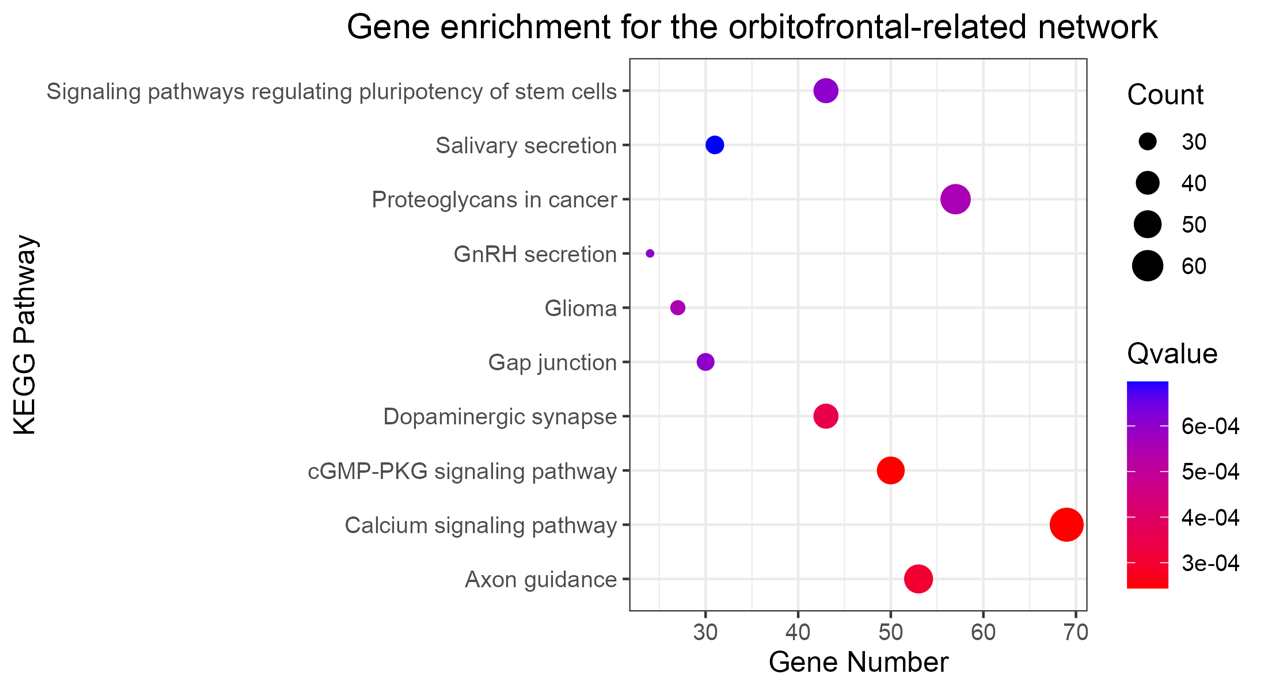

Figure S4 gene enrichment analysis for the orbitofrontal-related network

The top 10 significant GO terms and KEGG pathways were reported. The orbitofrontal-related network was associated with the dopaminergic synapse in the KEGG pathway analysis.

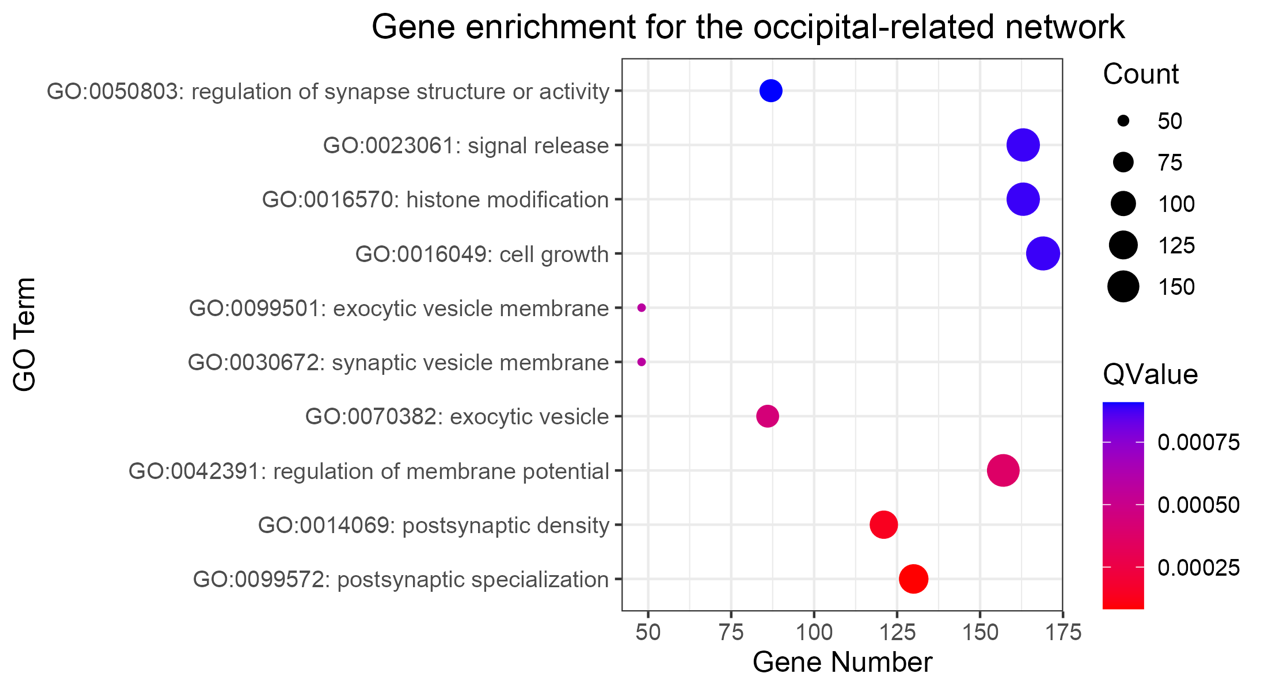

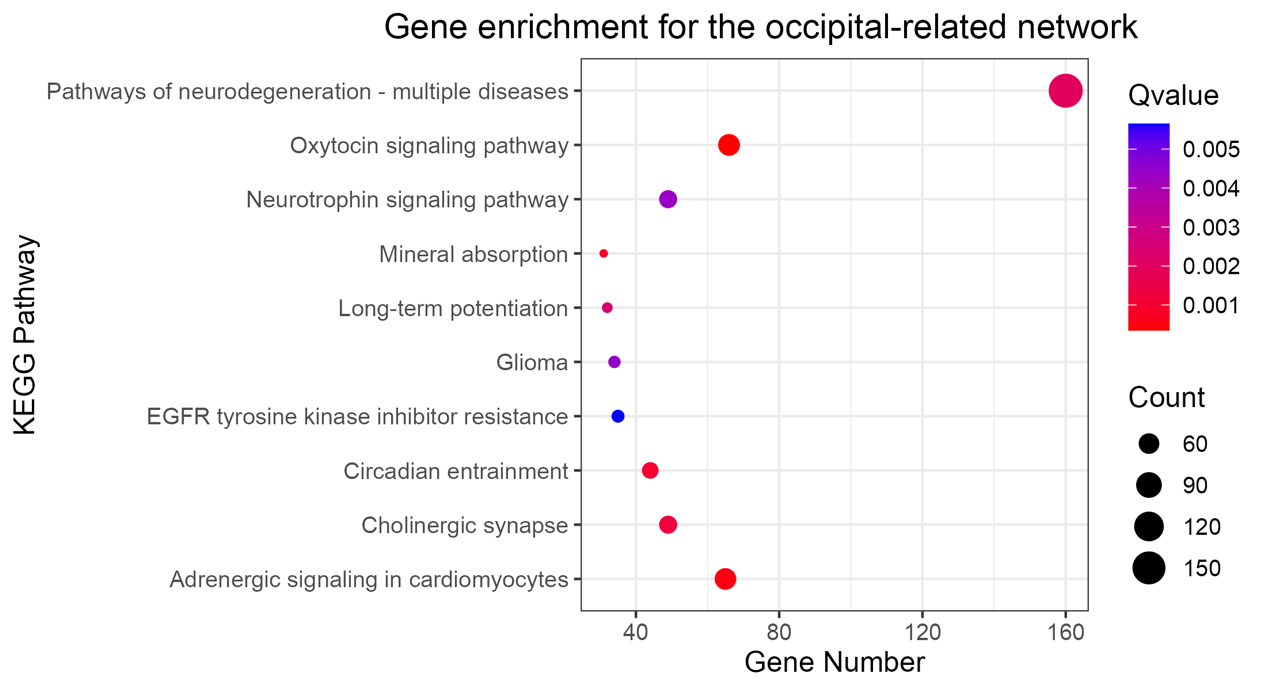

Figure S5 gene enrichment analysis for the occipital-related network

The top 10 significant GO terms and KEGG pathways were reported. The occipital-related network was not associated with the dopaminergic synapse in the KEGG pathway analysis.

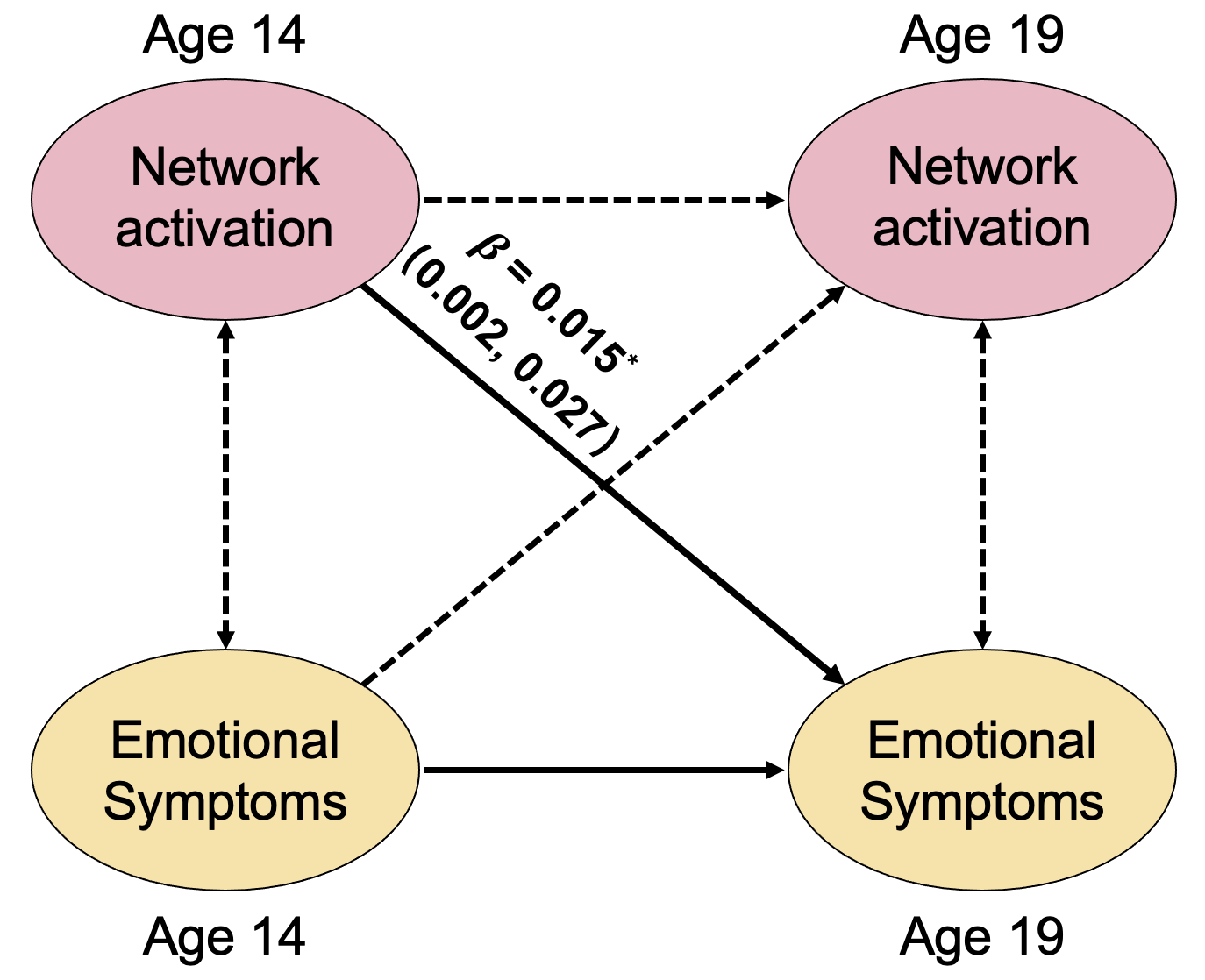

Figure S6 The cross-lagged panel model

After adjusting for both childhood abuse and PRS_MDD_, we found only one significant directionality in girls from the activation of the orbitofrontal-related network at age 14 to emotional symptoms at age 19, but not the other way around.

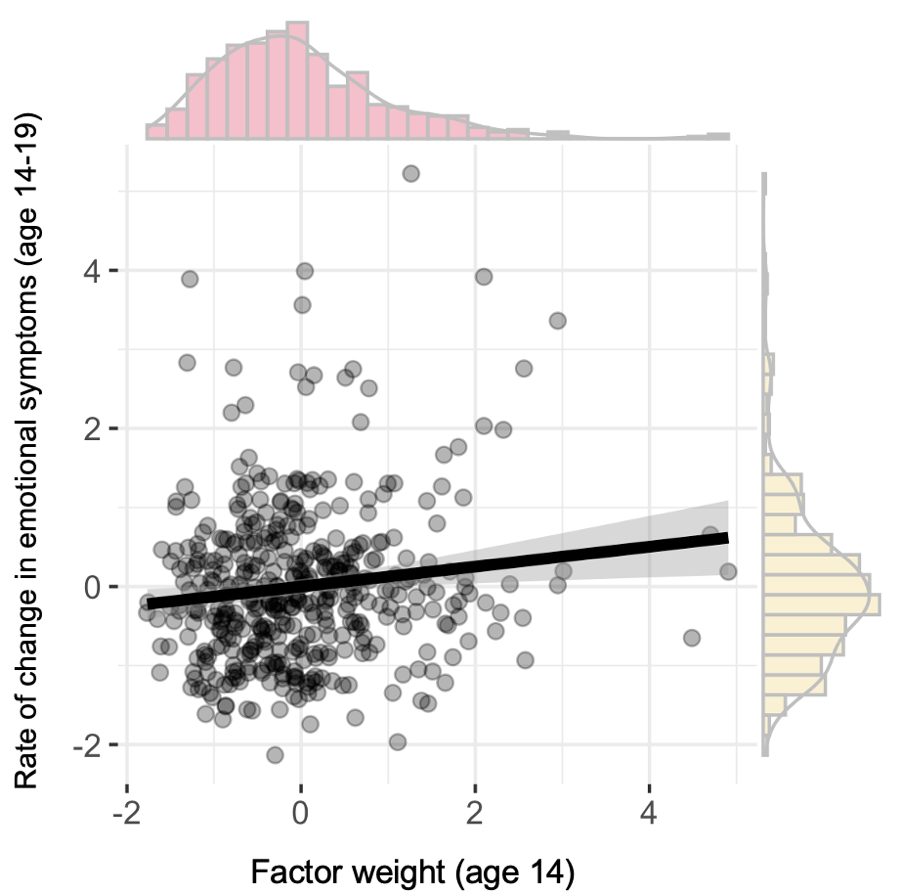

Figure S7 The directionality of the relationship between the orbitofrontal-related network and emotional symptoms in the cross-prediction model

Scatter plot. Residual scores were used after controlling for the covariates described in the Method S7. Factor weight, the activation of the orbitofrontal-related network.

### Supplementary Tables

Table S1 The main AAL2 regions of five latent networks

| **Networks** | **Regions** | **The number of voxels in AAL2** | **The number of voxels in this network** | **Proportion (voxel numbers of this network / voxel numbers of this region)** |
| --- | --- | --- | --- | --- |
| **Orbitofrontal-related network** | Frontal_Med_Orb_L | 219 | 219 | 100% |
|  | Frontal_Med_Orb_R | 269 | 264 | 98.14% |
|  | Frontal_Sup_Medial_L | 912 | 868 | 95.18% |
|  | Cingulate_Post_L | 134 | 127 | 94.78% |
|  | Cingulate_Ant_L | 411 | 379 | 92.21% |
|  | Angular_L | 335 | 305 | 91.04% |
|  | Cingulate_Ant_R | 391 | 348 | 89.00% |
|  | OFClat_L | 63 | 56 | 88.89% |
|  | Frontal_Sup_Medial_R | 579 | 499 | 86.18% |
|  | Olfactory_L | 79 | 68 | 86.08% |
|  | Angular_R | 481 | 377 | 78.38% |
|  | Cingulate_Post_R | 88 | 67 | 76.14% |
|  | OFClat_R | 66 | 48 | 72.73% |
|  | Frontal_Sup_2_L | 1434 | 1021 | 71.20% |
|  | Rectus_L | 255 | 179 | 70.20% |
|  | OFCpost_L | 183 | 126 | 68.85% |
|  | Frontal_Mid_2_L | 1329 | 910 | 68.47% |
|  | OFCant_R | 211 | 142 | 67.30% |
|  | Frontal_Inf_Orb_2_L | 228 | 150 | 65.79% |
|  | OFCant_L | 158 | 100 | 63.29% |
|  | Caudate_L | 289 | 182 | 62.98% |
|  | Rectus_R | 214 | 130 | 60.75% |
|  | Frontal_Sup_2_R | 1496 | 908 | 60.70% |
|  | ParaHippocampal_L | 297 | 175 | 58.92% |
|  | OFCpost_R | 173 | 100 | 57.80% |
|  | Frontal_Mid_2_R | 1452 | 753 | 51.86% |
|  | Olfactory_R | 79 | 39 | 49.37% |
|  | Parietal_Inf_R | 430 | 197 | 45.81% |
|  | Precuneus_L | 1051 | 478 | 45.48% |
|  | Hippocampus_L | 278 | 125 | 44.96% |
|  | ParaHippocampal_R | 315 | 134 | 42.54% |
|  | Caudate_R | 297 | 126 | 42.42% |
|  | Precuneus_R | 967 | 399 | 41.26% |
|  | Cingulate_Mid_L | 575 | 235 | 40.87% |
|  | Temporal_Mid_L | 1500 | 574 | 38.27% |
|  | Temporal_Pole_Mid_L | 226 | 84 | 37.17% |
|  | Frontal_Inf_Orb_2_R | 242 | 85 | 35.12% |
|  | Cingulate_Mid_R | 635 | 216 | 34.02% |
|  | Hippocampus_R | 292 | 98 | 33.56% |
| **Occipital-related network** | Lingual_R | 669 | 565 | 84.45% |
|  | Lingual_L | 661 | 557 | 84.27% |
|  | Cuneus_R | 457 | 377 | 82.49% |
|  | Heschl_R | 68 | 56 | 82.35% |
|  | Heschl_L | 77 | 61 | 79.22% |
|  | Parietal_Sup_R | 631 | 495 | 78.45% |
|  | Cuneus_L | 476 | 366 | 76.89% |
|  | Calcarine_L | 647 | 496 | 76.66% |
|  | Occipital_Sup_L | 409 | 301 | 73.59% |
|  | Rolandic_Oper_L | 297 | 207 | 69.70% |
|  | Insula_R | 515 | 358 | 69.51% |
|  | Parietal_Sup_L | 629 | 435 | 69.16% |
|  | Rolandic_Oper_R | 398 | 275 | 69.10% |
|  | Amygdala_L | 69 | 46 | 66.67% |
|  | Paracentral_Lobule_R | 235 | 153 | 65.11% |
|  | Calcarine_R | 525 | 341 | 64.95% |
|  | Occipital_Sup_R | 391 | 251 | 64.19% |
|  | Temporal_Sup_L | 667 | 414 | 62.07% |
|  | Fusiform_L | 693 | 402 | 58.01% |
|  | Fusiform_R | 756 | 371 | 49.07% |
|  | Temporal_Pole_Sup_L | 388 | 189 | 48.71% |
|  | Postcentral_L | 1186 | 574 | 48.40% |
|  | Paracentral_Lobule_L | 391 | 188 | 48.08% |
|  | Supp_Motor_Area_L | 648 | 308 | 47.53% |
|  | Postcentral_R | 1152 | 531 | 46.09% |
|  | Occipital_Mid_R | 607 | 265 | 43.66% |
|  | Amygdala_R | 69 | 30 | 43.48% |
|  | Precuneus_R | 967 | 418 | 43.23% |
|  | Insula_L | 536 | 228 | 42.54% |
|  | Precentral_R | 1000 | 425 | 42.50% |
|  | Supp_Motor_Area_R | 710 | 287 | 40.42% |
|  | Precuneus_L | 1051 | 414 | 39.39% |
|  | Temporal_Pole_Sup_R | 399 | 153 | 38.35% |
|  | Occipital_Inf_R | 329 | 125 | 37.99% |
|  | Cingulate_Mid_L | 575 | 207 | 36.00% |
|  | SupraMarginal_R | 537 | 192 | 35.75% |
|  | Frontal_Inf_Oper_R | 426 | 149 | 34.98% |
|  | Cingulate_Mid_R | 635 | 216 | 34.02% |
|  | Occipital_Inf_L | 269 | 91 | 33.83% |
|  | Parietal_Inf_R | 430 | 132 | 30.70% |
|  | Occipital_Mid_L | 966 | 291 | 30.12% |
| **Neutral>angry network 1** | Cuneus_L | 476 | 376 | 78.99% |
|  | Cingulate_Ant_L | 411 | 306 | 74.45% |
|  | Heschl_L | 77 | 57 | 74.03% |
|  | Cingulate_Post_L | 134 | 99 | 73.88% |
|  | Cingulate_Mid_L | 575 | 415 | 72.17% |
|  | Heschl_R | 68 | 48 | 70.59% |
|  | Precuneus_R | 967 | 644 | 66.60% |
|  | Precuneus_L | 1051 | 656 | 62.42% |
|  | Cingulate_Mid_R | 635 | 380 | 59.84% |
|  | Insula_R | 515 | 295 | 57.28% |
|  | Rolandic_Oper_L | 297 | 168 | 56.57% |
|  | Cuneus_R | 457 | 255 | 55.80% |
|  | Calcarine_L | 647 | 352 | 54.40% |
|  | Cingulate_Post_R | 88 | 45 | 51.14% |
|  | Frontal_Med_Orb_L | 219 | 111 | 50.68% |
|  | Rolandic_Oper_R | 398 | 199 | 50% |
|  | Frontal_Sup_Medial_L | 912 | 454 | 49.78% |
|  | Cingulate_Ant_R | 391 | 188 | 48.08% |
|  | Calcarine_R | 525 | 251 | 47.81% |
|  | Lingual_L | 661 | 309 | 46.75% |
|  | Temporal_Sup_L | 667 | 310 | 46.48% |
|  | Insula_L | 536 | 232 | 43.28% |
|  | Lingual_R | 669 | 271 | 40.51% |
|  | Frontal_Med_Orb_R | 269 | 90 | 33.46% |
| **Neutral>angry network 2** | OFCant_R | 211 | 205 | 97.16% |
|  | OFCant_L | 158 | 141 | 89.24% |
|  | Rectus_L | 255 | 223 | 87.45% |
|  | Rectus_R | 214 | 183 | 85.51% |
|  | Amygdala_L | 69 | 52 | 75.36% |
|  | OFCmed_L | 152 | 114 | 75.00% |
|  | Olfactory_R | 79 | 59 | 74.68% |
|  | Parietal_Sup_R | 631 | 456 | 72.27% |
|  | Olfactory_L | 79 | 57 | 72.15% |
|  | OFCpost_R | 173 | 123 | 71.10% |
|  | ParaHippocampal_L | 297 | 201 | 67.68% |
|  | OFCmed_R | 184 | 123 | 66.85% |
|  | Temporal_Pole_Mid_L | 226 | 150 | 66.37% |
|  | Paracentral_Lobule_R | 235 | 148 | 62.98% |
|  | OFCpost_L | 183 | 111 | 60.66% |
|  | ParaHippocampal_R | 315 | 191 | 60.63% |
|  | OFClat_R | 66 | 38 | 57.58% |
|  | Paracentral_Lobule_L | 391 | 225 | 57.54% |
|  | OFClat_L | 63 | 36 | 57.14% |
|  | Temporal_Inf_R | 1068 | 600 | 56.18% |
|  | Fusiform_L | 693 | 383 | 55.27% |
|  | Temporal_Inf_L | 932 | 514 | 55.15% |
|  | Parietal_Sup_L | 629 | 318 | 50.56% |
|  | Fusiform_R | 756 | 375 | 49.60% |
|  | Amygdala_R | 69 | 34 | 49.28% |
|  | Postcentral_R | 1152 | 438 | 38.02% |
|  | Frontal_Med_Orb_R | 269 | 91 | 33.83% |
|  | Temporal_Pole_Mid_R | 355 | 118 | 33.24% |
|  | Temporal_Pole_Sup_L | 388 | 126 | 32.47% |
|  | Occipital_Inf_L | 269 | 85 | 31.60% |
|  | Hippocampus_L | 278 | 87 | 31.29% |
|  | Occipital_Inf_R | 329 | 102 | 31.00% |
|  | Frontal_Med_Orb_L | 219 | 66 | 30.14% |
| **Neutral>angry network 3** | Frontal_Inf_Tri_R | 622 | 554 | 89.07% |
|  | Frontal_Inf_Tri_L | 733 | 652 | 88.95% |
|  | Frontal_Inf_Oper_R | 426 | 375 | 88.03% |
|  | Frontal_Inf_Oper_L | 315 | 264 | 83.81% |
|  | Frontal_Inf_Orb_2_L | 228 | 182 | 79.82% |
|  | Frontal_Inf_Orb_2_R | 242 | 186 | 76.86% |
|  | Temporal_Mid_R | 1310 | 859 | 65.57% |
|  | Temporal_Mid_L | 1500 | 922 | 61.47% |
|  | Temporal_Sup_R | 963 | 529 | 54.93% |
|  | SupraMarginal_L | 370 | 173 | 46.76% |
|  | SupraMarginal_R | 537 | 234 | 43.58% |
|  | Temporal_Pole_Sup_R | 399 | 160 | 40.10% |
|  | Occipital_Inf_R | 329 | 125 | 37.99% |
|  | Occipital_Inf_L | 269 | 98 | 36.43% |
|  | Precentral_L | 1044 | 374 | 35.82% |
|  | Frontal_Sup_Medial_R | 579 | 188 | 32.47% |
|  | Occipital_Mid_L | 966 | 309 | 31.99% |
|  | Temporal_Pole_Sup_L | 388 | 124 | 31.96% |

According to the voxel proportion of AAL2 regions for each network, the top 70% brain regions that are involved for each network are presented.

Table S2 Sex differences in weights of five latent networks

| Network | beta | t-value | p-value | 95%CI | Significant |
| --- | --- | --- | --- | --- | --- |
| Orbitofrontal-related network | -0.081 | -1.133 | 0.258 | [-0.221, 0.059] | NS |
| Occipital-related network | -0.23 | -3.228 | 0.001 | [-0.369, -0.09] | ** |
| Neutral>angry network 1 | -0.142 | -2.008 | 0.045 | [-0.282, -0.003] | * |
| Neutral>angry network 2 | -0.071 | -0.991 | 0.322 | [-0.211, 0.069] | NS |
| Neutral>angry network 3 | 0.104 | 1.481 | 0.139 | [-0.034, 0.243] | NS |

Table S3 Developmental trajectories of latent networks from age 14 to 19 years in boys

| Network | F value | Sum Square | p value | Mean±SD (age 14) | Mean±SD (age 19) | Trend |
| --- | --- | --- | --- | --- | --- | --- |
| Orbitofrontal-related network | 4.509 | 1401 | **0.034** | 26.996±18.252 | 29.714±19.865 | increase |
| Occipital-related network | 4.593 | 2123 | **0.033** | 26.958±19.685 | 30.305±24.120 | increase |
| Neutral>angry network 1 | 2.021 | 438 | 0.156 | 18.070±12.882 | 16.549±16.034 |  |
| Neutral>angry network 2 | 0.558 | 114 | 0.456 | 18.843±14.397 | 18.068±15.614 |  |
| Neutral>angry network 3 | 5.356 | 968 | **0.021** | 20.087±12.386 | 22.347±15.801 | increase |

Table S4 Developmental trajectories of latent networks from age 14 to 19 years in girls

| Network | F value | Sum Square | p value | Mean±SD (age 14) | Mean±SD (age 19) | Trend |
| --- | --- | --- | --- | --- | --- | --- |
| Orbitofrontal-related network | 4.142 | 860 | **0.042** | 26.163±15.272 | 28.170±15.612 | increase |
| Occipital-related network | 0.215 | 56 | 0.643 | 24.988±16.840 | 25.497±16.599 |  |
| Neutral>angry network 1 | 8.097 | 1277 | **0.004** | 16.653±12.352 | 14.215±12.679 | decrease |
| Neutral>angry network 2 | 1.938 | 282 | 0.165 | 18.021±12.049 | 16.876±12.602 |  |
| Neutral>angry network 3 | 4.053 | 587 | **0.045** | 22.104±12.347 | 23.758±13.441 | increase |

Table S5 Three-way interactions among childhood abuse, PRS­_MDD_ and the activation of each latent networks in relation to emotional symptoms in girls at age 19

| Interaction | beta | 95%CI | t | p-value | FDR-p |
| --- | --- | --- | --- | --- | --- |
| Orb x CA x PRS_MDD_ | -0.128 | [-0.224, -0.031] | -2.594 | 0.009 | **0.023** |
| Occ x CA x PRS_MDD_ | -0.148 | [-0.253, -0.043] | -2.777 | 0.005 | **0.023** |
| NA1 x CA x PRS_MDD_ | 0.182 | [0.021, 0.343] | 2.226 | 0.026 | **0.043** |
| NA2 x CA x PRS_MDD_ | 0.002 | [-0.182, 0.186] | 0.017 | 0.986 | 0.986 |
| NA3 x CA x PRS_MDD_ | -0.058 | [-0.208, 0.092] | -0.763 | 0.446 | 0.558 |

Orb, the activation of the orbitofrontal-related network; Occ, the activation of the occipital-related network; NA1, the activation of the neutral>angry network 1; NA2, the activation of the neutral>angry network 2; NA3, the activation of the neutral>angry network 3; CA, childhood abuse; PRS_MDD_, Polygenic risk scores for major depression disorder.

Table S6 Three-way interaction effects after binarizing the childhood abuse

| Interaction | beta | 95%CI | t | p-value |
| --- | --- | --- | --- | --- |
| Orb x CA x PRS_MDD_ | -0.299 | [-0.516, -0.081] | -2.699 | **0.007** |
| Occ x CA x PRS_MDD_ | -0.252 | [-0.511, 0.006] | -1.922 | 0.055 |
| NA1 x CA x PRS_MDD_ | 0.291 | [-0.032, 0.616] | 1.773 | 0.077 |

Childhood abuse was binarized according to the clinical cut-offs (See Methods). Orb, the activation of the orbitofrontal-related network; Occ, the activation of the occipital-related network; NA1, the activation of the neutral>angry network 1; CA, childhood abuse; PRS_MDD_, Polygenic risk scores for major depression disorder.

Table S7 Three-way interaction effects controlling for additional covariates in girls

| Covariates | Interaction | beta | 95%CI | t | p-value |
| --- | --- | --- | --- | --- | --- |
| Age | Orb x CA x PRS_MDD_ | -0.124 | [-0.221, -0.027] | -2.508 | **0.012** |
|  | Occ x CA x PRS_MDD_ | -0.145 | [-0.250, -0.040] | -2.704 | **0.007** |
|  | NA1 x CA x PRS_MDD_ | 0.176 | [0.014, 0.337] | 2.134 | **0.033** |
| Covariates | Interaction | beta | 95%CI | t | p-value |
| Childhood neglect | Orb x CA x PRS_MDD_ | -0.148 | [-0.245, -0.053] | -3.045 | **0.002** |
|  | Occ x CA x PRS_MDD_ | -0.164 | [-0.268, -0.060] | -3.116 | **0.001** |
|  | NA1 x CA x PRS_MDD_ | 0.180 | [0.021, 0.340] | 2.229 | **0.026** |
| Covariates | Interaction | beta | 95%CI | t | p-value |
| IQ | Orb x CA x PRS_MDD_ | -0.132 | [-0.229, -0.035] | -2.672 | **0.008** |
|  | Occ x CA x PRS_MDD_ | -0.136 | [-0.243, -0.029] | -2.504 | **0.013** |
|  | NA1 x CA x PRS_MDD_ | 0.171 | [0.012, 0.330] | 2.110 | **0.035** |
| Covariates | Interaction | beta | 95%CI | t | p-value |
| Smoking | Orb x CA x PRS_MDD_ | -0.127 | [-0.224, -0.030] | -2.573 | **0.010** |
|  | Occ x CA x PRS_MDD_ | -0.148 | [-0.253, -0.043] | -2.769 | **0.006** |
|  | NA1 x CA x PRS_MDD_ | 0.180 | [0.019, 0.342] | 2.201 | **0.028** |
| Covariates | Interaction | beta | 95%CI | t | p-value |
| Alcohol use | Orb x CA x PRS_MDD_ | -0.127 | [-0.224, -0.031] | -2.586 | **0.010** |
|  | Occ x CA x PRS_MDD_ | -0.148 | [-0.253, -0.043] | -2.778 | **0.006** |
|  | NA1 x CA x PRS_MDD_ | 0.183 | [0.022, 0.344] | 2.230 | **0.026** |
| Covariates | Interaction | beta | 95%CI | t | p-value |
| Drug use | Orb x CA x PRS_MDD_ | -0.122 | [-0.219, -0.025] | -2.484 | **0.013** |
|  | Occ x CA x PRS_MDD_ | -0.144 | [-0.249, -0.039] | -2.695 | **0.007** |
|  | NA1 x CA x PRS_MDD_ | 0.172 | [0.011, 0.333] | 2.099 | **0.036** |

Orb, the activation of the orbitofrontal-related network; Occ, the activation of the occipital-related network; NA1, the activation of the neutral>angry network 1; CA, childhood abuse; PRS_MDD_, Polygenic risk scores for major depression disorder.

Table S8 Three-way interaction effects showing specificity to emotional symptoms in girls

| The association between childhood abuse and the other four dimensions in the SDQ | | | | | |
| --- | --- | --- | --- | --- | --- |
| Predictor | Outcome | beta | 95%CI | t | p-value |
| CA | Conduct | 0.175 | [0.095, 0.255] | 4.294 | **<0.001** |
|  | Hyper | 0.101 | [0.020, 0.182] | 2.459 | **0.014** |
|  | Peer | 0.104 | [0.024, 0.184] | 2.550 | **0.011** |
|  | Prosocial | -0.047 | [-0.116, 0.022] | -1.335 | 0.182 |
| Replacing the emotional symptom score with the other four dimensions of behavioral problem scores in the SDQ | | | | | |
| Formula | Interaction | beta | 95%CI | t | p-value |
| Conduct~W:CA: PRS_MDD_ | Orb x CA x PRS_MDD_ | -0.052 | [-0.147, 0.044] | -1.062 | 0.289 |
|  | Occ x CA x PRS_MDD_ | 0.041 | [-0.063, 0.145] | 0.773 | 0.440 |
|  | NA1x CA x PRS_MDD_ | -0.061 | [-0.220, 0.098] | -0.753 | 0.452 |
| Formula | Interaction | beta | 95%CI | t | p-value |
| Hyper~W:CA: PRS_MDD_ | Orb x CA x PRS_MDD_ | -0.076 | [-0.172, 0.021] | -1.539 | 0.124 |
|  | Occ x CA x PRS_MDD_ | -0.104 | [-0.209, 0.0003] | -1.959 | 0.051 |
|  | NA1 x CA x PRS_MDD_ | -0.022 | [-0.183, 0.139] | -0.271 | 0.791 |
| Formula | Interaction | beta | 95%CI | t | p-value |
| Peer~W:CA: PRS_MDD_ | Orb x CA x PRS_MDD_ | -0.083 | [-0.181, 0.014] | -1.670 | 0.096 |
|  | Occ x CA x PRS_MDD_ | -0.054 | [-0.161, 0.052] | -1.003 | 0.316 |
|  | NA1 x CA x PRS_MDD_ | 0.078 | [-0.085, 0.241] | 0.934 | 0.351 |
| Formula | Interaction | beta | 95%CI | t | p-value |
| Prosocial~W:CA: PRS_MDD_ | Orb x CA x PRS_MDD_ | 0.085 | [-0.010, 0.179] | 1.753 | 0.080 |
|  | Occ x CA x PRS_MDD_ | 0.138 | [-0.023, 0.235] | 1.631 | 0.078 |
|  | NA1 x CA x PRS_MDD_ | -0.172 | [-0.278, 0.012] | -1.151 | 0.082 |
| Replacing the PRS_MDD_ with the PRS_ADHD_ or the PRS_SCZ_ | | | | | |
| Modulation | Interaction | beta | 95%CI | t-value | p-value |
| PRS_ADHD_ | Orb$\times$CA$\times$PRS_ADHD_ | 0.025 | [-0.081, 0.131] | 0.471 | 0.638 |
|  | Occ$\times$CA$\times$PRS_ADHD_ | 0.028 | [-0.098, 0.154] | 0.064 | 0.664 |
| PRS_SCZ_ | Orb$\times$CA$\times$PRS_SCZ_ | -0.049 | [-0.150, 0.053] | -0.943 | 0.346 |
|  | Occ$\times$CA$\times$PRS_SCZ_ | -0.040 | [-0.164, 0.084] | -0.638 | 0.524 |

Conduct, conduct symptoms; Hyper, hyperactivity/inattention symptoms; Peer, peer relationship problems; Prosocial, prosocial behavior; W, network activations; CA, childhood abuse; PRS_MDD,_ Polygenic risk scores for major depression disorder; Orb, the activation of the orbitofrontal-related network; Occ, the activation of the occipital-related network; NA1, the activation of the neutral>angry network 1.

Table S9 Longitudinal relationship between the development of latent networks and the development of emotional symptoms in girls

| **From network activation at age 14 to the rate of change in the emotional symptom score** | | | | | |
| --- | --- | --- | --- | --- | --- |
| Predictor | beta | 95%CI | t | p-value | FDR-p |
| Orb | 0.128 | [0.029, 0.227] | 2.542 | 0.011 | **0.033** |
| Occ | -0.004 | [-0.100, 0.092] | -0.078 | 0.938 | 0.938 |
| NA1 | -0.064 | [-0.160, 0.031] | -1.321 | 0.187 | 0.281 |
| **From emotional symptom score at age 14 to the rate of change in the activations of three networks** | | | | | |
| Regressor | beta | 95%CI | t | p-value |  |
| Orb | 0.003 | [-0.095, 0.101] | 0.062 | 0.950 |  |
| Occ | -0.003 | [-0.084, 0.075] | -0.085 | 0.932 |  |
| NA1 | -0.007 | [-0.086, 0.068] | -0.176 | 0.861 |  |

Orb, the activation of the orbitofrontal-related network; Occ, the activation of the occipital-related network; NA1, the activation of the neutral>angry network 1.
